# Genetic diversity of the pathogenic banana Fusarium wilt in northern Viet Nam

**DOI:** 10.1101/2021.05.26.445774

**Authors:** Loan Le Thi, Arne Mertens, Dang Toan Vu, Dang Tuong Vu, Pham Le Anh Minh, Huy Nguyen Duc, Sander de Backer, Rony Swennen, Filip Vandelook, Bart Panis, Mario Amalfi, Cony Decock, Sofia I.F. Gomes, Vincent S.F.T. Merckx, Steven B. Janssens

## Abstract

*Fusarium* is one of the most important fungal genera of plant pathogens that affect the cultivation of a wide range of crops. Agricultural losses caused by *Fusarium oxysporum* f. sp. *cubense (Foc)* have a direct effect on the income, subsistence and nourishment of thousands of smallholder farmers worldwide. In addition, also commercial growers are strongly affected. For Viet Nam, predictions on the impact of *Foc* for the future are dramatic with an estimated loss in banana production area of 8% within the next 5 years and up to 71% within the next 25 years. In the current study we applied a combined morphological - molecular approach to assess the taxonomic identity and phylogenetic position of the different *Foc* isolates that were collected in northern Viet Nam. In addition, we aim to estimate the proportion of the different *Fusarium* races that are infecting bananas in northern Viet Nam. The morphology of the isolates was investigated by growing the collected *Fusarium* isolates on four distinct nutritious media (PDA, SNA, CLA, and OMA). Molecular phylogenetic relationships were inferred by sequencing partial *rpb1*, *rpb2* and *tef1a* genes and adding the obtained sequences into a large phylogenetic framework. The present study showed that *Foc* Race 1 is the most common strain in northern Viet Nam, causing 74% of all the infections. A more in-depth molecular characterization shows that of the *Foc* Race 1 infections, 92% are caused by *Fusarium tardichlamydosporum* and 8% by *F. duoseptatum*. Compared to *Foc* Race 1, *Foc* TR4 (represented by *F. odoratissimum*) account for only 10.5% of the Fusarium wilting in northern Viet Nam demonstrating that *Foc* TR4 is not yet a dominant strain in the region. *Foc* Race 2 infections of Vietnamese bananas also account for 10.5% of the Fusarium wilting in northern Viet Nam. One of the isolates cultured from diseased bananas collected in northern Viet Nam was phylogenetically not positioned within the *F. oxysporum* species complex (FOSC), but in contrast fell within the *Fusarium fujikuroi* species complex (FFSC). As a result, it is possible that a new pathogen for bananas has been found. Besides being present on several ABB ‘Tay banana’, *Foc* Race 1 was also found as pathogen on wild *Musa lutea*, showing the importance of wild bananas as possible sink for *Foc*.

## Introduction

The ascomycete genus *Fusarium* comprises one of the most important fungal plant pathogens impacting the cultivation of numerous agricultural crops (e.g., rice, coffee, tomato, melon, wheat; Dean et al. 2012). *Fusarium* has a considerable economic, social and biological impact on the daily livelihood of millions of people worldwide. *Fusarium oxysporum* is one of the two most devastating pathogens in the genus, besides *F. gramineum*. The *Fusarium oxysporum* species complex is responsible for wilt diseases of various crops (e.g. cotton and tomato wilt), but is mainly known from its massive impact on the banana industry (Panama disease). For more than 100 years, the fungus has affected banana production worldwide (Ploetz and Pegg 1997; Ploetz 2015a). For millions of people, bananas are an important food crop. With an annual global production of 153 million tons produced on 5.6 million hectares of land, a revenue of more than 26.5 billion Euro was generated in 2017 (FAO 2018). Particularly in Asia, Africa, Latin America and the Caribbean bananas support rural livelihood as most of the grown bananas are self-consumed or locally traded. As a result, any reduction in crop harvest caused by for example *Foc* infections has a direct effect on the income, subsistence and nourishment of thousands of smallholders. Additionally, also the worldwide banana export is seriously affected by *Foc*, as most of its current production depends on the cultivation of members of the Cavendish subgroup (Buddenhagen 2009, Ploetz 2009). Although these triploid ‘AAA’ Cavendish cultivars were selected in the past century for their resistance against *F. oxysporum* f. sp. *cubense* Race 1 (*Foc*-Race 1), to which the initially grown Gros Michel cultivars were highly susceptible (Stover 1962a), Cavendish cultivars (e.g., Grand Naine, Williams) are highly susceptible to *Foc*-TR4. All *Foc* strains currently known (e.g., Race 1, Race 2, STR4, TR4) pose a huge treat for banana cultivation worldwide. Moreover, knowing that nearly half of the global banana production is derived from Cavendish clones, and that they also become more popular for domestic use, a *Foc*-TR4 pandemic is still not averted to date (Ploetz 2015b; Zheng et al. 2018).

In the near future, *Foc* will further intensively spread in Asia, thereby significantly affecting important banana producing countries such as China, the Philippines, Pakistan and Viet Nam (Scheerer et al. 2018). For Viet Nam, the predictions are dramatic, estimating a loss in banana production area for the country of 8% within the next 5 years and up to 71% within the next 25 years (Scheerer et al. 2018).

As a soil-borne fungus, *Foc* invades the rooting system from where it subsequently moves into the vascular tissue that gradually deteriorates. When reaching the corm, wilting takes place eventually resulting in the death of the contaminated plant (Stover 1962a). A particular problem that arises with *Foc* infections is the remaining presence of *Foc* spores (microconidia, macroconidia and chlamydospores) in the soil surrounding the infected plants for at least 20 years after the complete removal of all infected plants or plant tissue (Stover 1962b; Buddenhagen 2009; Dita et al. 2018). As a result, reinfection of new banana accessions in the same area is very likely to happen in the absence of a complete soil disinfection or if one has not waited long enough for planting new *Musa* cultivars (Moore et al. 2001, Huang et al. 2012). Therefore, *Fusarium* wilt not only has an impact on the overall yield during the time of infection, but also on the land use for banana cultivars during the coming 20 years. In 1968, Vakili and coworkers published a first survey on the presence of *Fusarium* infected bananas in Southern Viet Nam (Vakili et al. 1968). Later studies showed that by the end of the 20^th^ century, *Foc* infections were omnipresent in the whole country (Mai Van Tri 1997; Bentley et al. 1998; Vinh et al. 2001). Characterization of the *Fusarium* isolates in the above-mentioned studies demonstrated that *Fusarium* wilting on bananas in Viet Nam was derived from different *Foc* strains (e.g., VCG 0123, VCG 0124, VCG 0124/5, VCG 0125). Hung et al. (2017) reported the first observation of *Foc*-TR4 (VCG 01213/16) on Cavendish bananas in Viet Nam using a combined molecular (PCR approach) and morphological characterization. However, Zheng et al. (2018) claimed that they made the earliest collected records of *Foc*-TR4 in Viet Nam in 2016, by assessing the pathogenicity of the collected strains and characterizing them molecularly using whole genome sequencing methodology. The study of Mostert et al. (2017) also made use of a molecular-morphological characterization approach to determine the origin of the different *Fusarium* infections in Viet Nam. Their results showed the presence of at least five different *Foc* strains (VCG 0123, VCG 0124, VCG 0124/5, VCG 0128, VCG 01221) of which the latter two were not yet detected in earlier studies.

Whereas pathogenic *Foc* lineages were usually classified into three races (*Foc* 1, 2 & 4) based on the different *Musa* cultivars they had infected, the development of the Vegetative Compatibility Group (VCG) system resulted in a more in-depth identification tool of *Foc* strains into 24 unique entities (Fourie et al. 2011; O’Donnell et al. 2009; Perez-Vicente et al. 2014; Mostert et al. 2017). The fact that isolated *Foc* lineages could already be split up in compatible vegetative groups already indicated that there are more natural lineages in the FOSC (*Fusarium oxysporum* species complex) than can be reflected by the number of races. In addition, the polyphyletic nature of *Fusarium oxysporum* f. sp. *cubense* isolates is also demonstrated by Maryani et al. (2019a), who used a combined molecular phylogenetic approach to delineate natural lineages within the *Fusarium oxysporum* species complex (FOSC; O’Donnell and Cigelnik 1997), thereby describing 11 new *Fusarium* species which were formerly considered as *Fusarium oxysporum*. A side result of the study of Maryani et al. (2019) also indicated that the VCG system is perhaps slightly prone to an oversimplification of the categorization of different *Foc* strains that cause *Fusarium* wilting in bananas and plantains. As a result, in the current study we aim to assess the overall diversity of *Foc* wilting in northern Viet Nam by using combined morphological - molecular phylogenetic approach in which also the different VCG’s are included. With this approach we provide the overall species identity and phylogenetic position of *Foc* infections in the northern Vietnamese region and examine the genetic diversity between the different *Foc* isolates (from wild and cultivated bananes) that were collected from various provinces in northern Viet Nam. Furthermore, our results will give an indication of the proportion of the different *Foc* strains (and linked VCG’s) that are currently infecting bananas in northern Viet Nam.

## Material and methods

### Sampling

From April 2018 until December 2019, several field trips were carried out focusing on the presence of banana *Fusarium* wilt in northern Viet Nam. During these surveys, banana *Fusarium* wilt samples were collected at 19 locations situated in three large geographic regions (North-eastern region, Red River Delta and North Central region; Table 1, Fig. 1). *Fusarium* infected banana plants were identified by following a set of diagnostic characters in which (mostly older) leaves were clearly yellow (initiated from the leaf margin) or even completely collapsed, halfway the petiole forming a ring of dead leaves around a dying plant, combined with brown discoloration and longitudinally fissuring of the pseudostems leaf sheaths (Fig. 2). From symptomatic plants observed in the field, discoloured brownish vascular tissue was collected from pseudostems and roots. Subsequent to collection, infected tissue samples were stored in paper bags and put in a refrigerator or cooling box to avoid quality loss upon further analysis in the molecular lab. For each sample collected, notes were taken about the altitude, longitude and latitude, site location and the host specimen. Collected *Fusarium* samples were stored at the Plant Resources Center (PRC), Ha Noi, Viet Nam. Of the 19 collected *Fusarium* samples, 17 were found in the triploid *Musa* ABB variety ‘Tay banana’, one in the triploid *Musa* AAA variety ‘Cavendish’, and one on a wild *Musa lutea* specimen (Table 1).

**Fig. 1.**
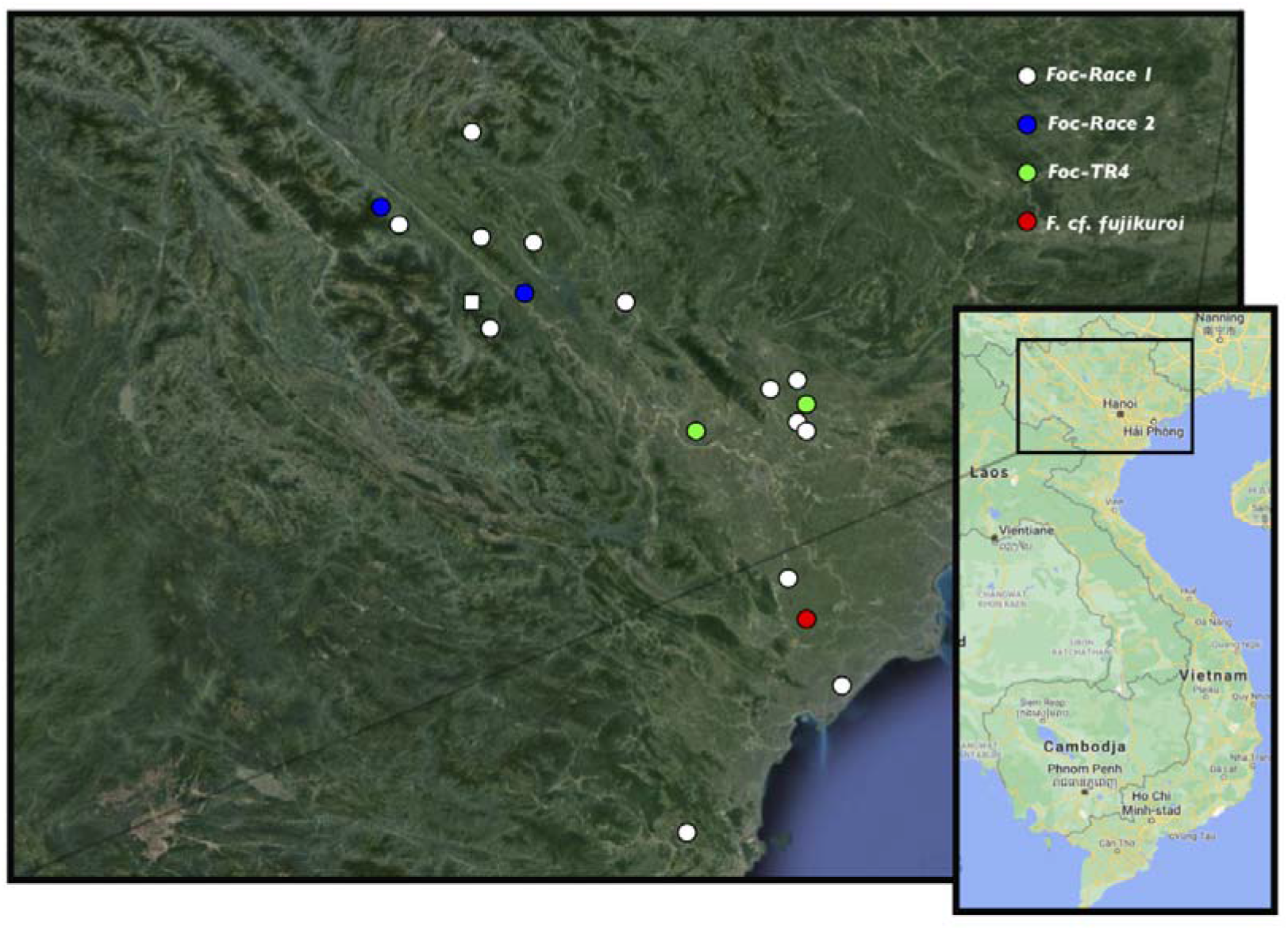
Distribution map of localities in northern Viet Nam where Fusarium wilting was observed. Colours indicate different *Fusarium* strains or species. Squares indicate *Fusarium* infections of wild bananas; circles indicate infections of cultivated bananas.

**Fig. 2.**
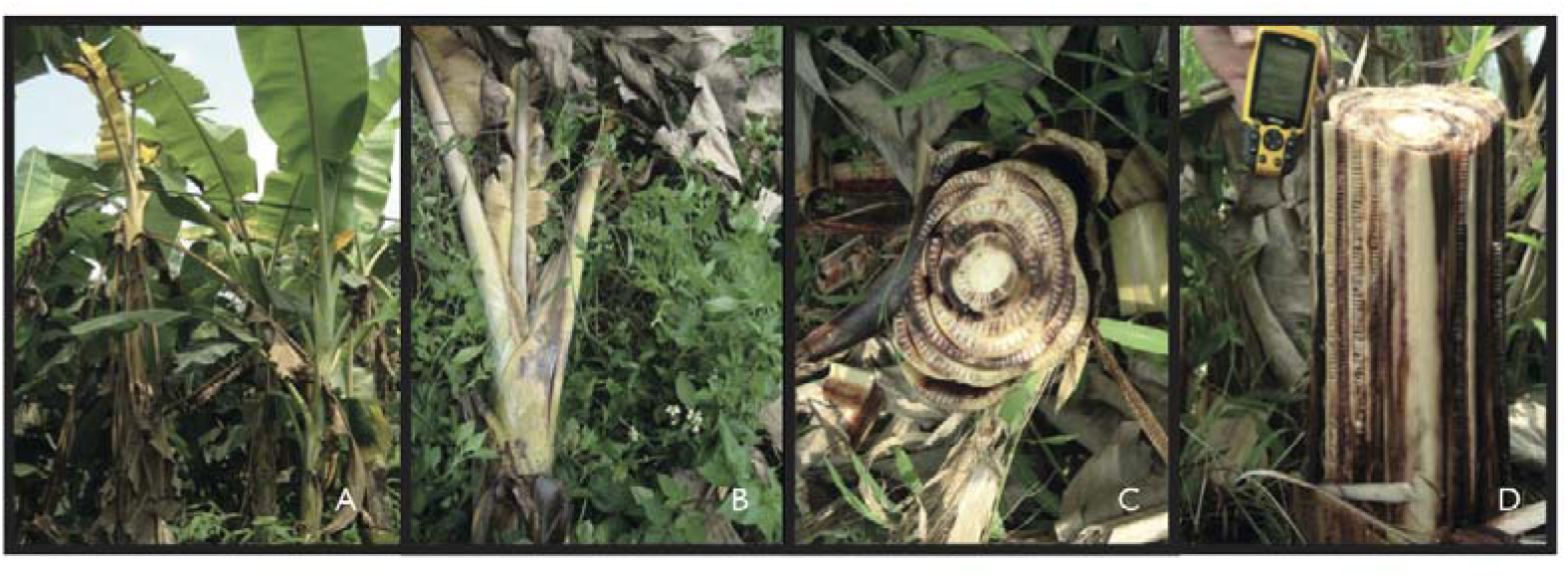
A. Overall view of a banana plant infected by *Fusarium* wilt, B. Detailed view of wilted plant. C. Radial cutting of *Fusarium* infected banana pseudostem. D. Tangential cutting of *Fusarium* infected banana pseudostem.

**Table 1.** List of collected *Fusarium* (*Foc*) wilt samples on bananas in northern Viet Nam

| Isolate | Locality | Cultivar or Species | Altitude (m) | Latitude | Longitude |
| --- | --- | --- | --- | --- | --- |
| FOC1 | Yen Binh town, Quang Binh district, Ha Giang province | Tay banana (ABB) | 78 | 22°23'5.28" | 104°32'54.0" |
| FOC2 | Dong Cay village, Yen Thang commune, Luc Yen district, Yen Bai province | Tay banana (ABB) | 188 | 22°05'46.2" | 104°45'34.0" |
| FOC4 | Khe Chao village, Ngoi A commune, Van Yen district, Yen Bai province | Tay banana (ABB) | 76 | 21°54'21.4" | 104°43'48.3" |
| FOC5 | No 7 village, Dai Son commune, Van Yen district, Yen Bai province | *Musa lutea* | 351 | 21°48'17.6" | 104°36'01.0" |
| FOC6-1 | No 4 village, Dai Son commune, Van Yen district, Yen Bai province | Tay banana (ABB) | 352 | 21°48'16.9" | 104°36'02.5" |
| FOC7 | No 18 village, Lam Giang commune, Van Yen district, Yen Bai province | Tay banana (ABB) | 137 | 22°02'35.3" | 104°30'52.3" |
| FOC10 | Hai Son 1 village, Phu Nhuan commune, Bao Thang district, Lao Cai province | Tay banana (ABB) | 370 | 22°13'58.6" | 104°08'14.9" |
| FOC11 | Khanh Yen town, Van Ban district, Lao Cai province | Tay banana (ABB) | 383 | 22°06'53.9" | 104°14'37.9" |
| FOC16 | Hoang An commune, Hiep Hoa district, Bac Giang province | Tay banana (ABB) | 20 | 21°23'00.2" | 105°58'40.5" |
| FOC18 | Hoang Thanh commune, Hiep Hoa district, Bac Giang province | Tay banana (ABB) | 18 | 21°23'22.1" | 106°00'24.9" |
| FOC21 | Thinh Long town, Hai Hau district, Nam Dinh province | Tay banana (ABB) | 4 | 20°01'59" | 106°12'49.0" |
| FOC23-2 | Thanh Chau commune, Phu Ly district, Ha Nam province | Tay banana (ABB) | 7 | 20°31'24.2" | 105°55'31.1" |
| FOC24 | Hung Thanh commune, Tuyen Quang city, Tuyen Quang province | Tay banana (ABB) | 26 | 21°48'19.4" | 105°11'39.7" |
| FOC25-1 | Le Chi commune, Gia Lam district, Ha Noi province | Tay banana (ABB) | 11 | 21°18'20.5" | 106°00'24.6" |
| FOC25-2 | Le Chi commune, Gia Lam district, Ha Noi province | Tay banana (ABB) | 11 | 21°18'20.5" | 106°00'24.6" |
| FOC38 | Quan Mia village, Nghia Tan commune, Nghia Dan district, Nghe An province | Tay banana (ABB) | 85 | 19°19'07.5" | 105°21'53.7" |
| FOC56 | Agriculture University, Trau Quy town, Gia Lam district, Ha Noi province | Tay banana (ABB) | 11 | 21°18'20.5" | 106°00'24.6" |
| FOC58 | Nui Ngam, Minh Tan commune, Vu Ban district, Nam Dinh province | Tay banana (ABB) | 3 | 20°21'59.1"N | 106°04'02.9"E |
| FOC 61 | Hong Chau commune, Yen Lac district, Vinh Phuc province | Cavendish (AAA) | 3 | 21°10'17.1"N | 105°34'37.0"E |

### Isolate cultivation

In order to observe possible morphological differences between the *Fusarium* wilt isolates that were collected from the wild and cultivated northern Vietnamese *Musa* accessions, we followed the approach of Groenewald et al. (2006) in which different *Foc* isolate were grown on different growing media. All 19 *Fusarium* isolates were cultured prior to further analysis.

Infected discoloured pseudostem tissue samples were cut into 2-3 cm pieces and placed on the Komada medium (Komada 1975). After a few days, fungal *Fusarium* colonies were transferred to plates with different medium and were then put in a growing chamber at 25^°^ C until the colonies reached a size of 2-3 cm. The different isolates were grown on four distinct nutritious media to observe the *Fusarium* wilting in different culture conditions: PDA (Potato Dextrose Agar), SNA (Spezieller Nährstoffarmer Agar), CLA (Carnation Leaf Agar), and OMA (Oatmeal Agar)(Nirenberg 1976). The PDA medium consisted of 200g potato dextrose, 20g D-glucose and 20g agar dissolved in 1000ml distilled water, whereas the SNA medium consisted of 1g KH_2_PO_4_, 1g KNO_3_, 0.5g MgSO_4_•7H_2_O, 0.5g KCl, 0.2g D-glucose and 0.2g D-sucrose dissolved in 1000ml distilled water. The CLA medium contained aseptic carnation leaves and 20g agar dissolved in 1000ml distilled water. The OMA medium consisted of 50g oatmeal and 20g agar dissolved in 1000ml distilled water. Growth of the *Fusarium* isolates on the different media took place under in-vitro conditions with the optimal grow temperature between 23 and 27°C (Pérez et al. 2003). After 7 days of incubation, the developing colonies were morphologically investigated under a light microscope (400x magnification). Coloration of the colony, the morphology and size of the conidia were determined. The colony reverse colour was determined on PDA medium after incubation at room temperature. Colony colours were assessed with the colour charts of Rayner (1970). In addition to colony colour, also the aroma of the different cultures was assessed as a strong rank odour generated by mature cultures is a typical characteristic for TR4 infections. In the first stage of culturing, we characterized the isolates as *Fusarium* spp. emanated from mycelium morphology and the presence of different types of conidia. The study of Maryani et al. (2019a) was used to further classify the Foc lineages into different sublineages. All obtained *Fusarium* isolates were stored in the Plant Resources Center (PRC), Ha Noi, Viet Nam.

### Molecular protocols

In order to extract high-quality DNA from the *Fusarium* wilt isolates collected and cultured, we used the pure mycelium cultures that were generated for the morphological characterization of the banana wilt. Total genomic DNA was isolated using a modified TNE protocol based on the study of Lin et al. (2008) and Dellaporte et al. (1983). After the addition of 5ml TNE buffer (100 mM Tris-HCl, 50 mM EDTA, 50 mM NaCl, 8 μM β-mercaptoethanol, 1% SDS, pH 8.0) to the sampled mycelium, the samples were incubated for 1h (65°C). Subsequent to the lysis phase, 1.66ml NaOAc (5M) was added and centrifuged. Chloroform-isoamylalcohol (24/1 v/v) extraction was done twice, followed by an isopropanol precipitation at −32°C for 12h. After centrifugation at 4°C, the pellet was washed twice (75% ethanol), air-dried, and dissolved in 100µl TE buffer (10mM TrisHCl, 0.1mM EDTA; pH 8) Amplification reactions of *rpb1*, *rpb2* and *tef1a* were carried out using standard PCR (20µl). Reactions were initiated with a 3 min heating at 95°C followed by 30 cycles consisting of 95°C for 30s, 55-65°C (*rpb1* and *rpb2*) and 53°-59°C (*tef1a*) for 60s, and 72°C for 60s. Reactions ended with a 3 min incubation at 72°C. Primers designed by O’Donnell et al. (1998) were used to sequence *tef1a*, whereas primers for *rpb1* and *rpb2* were adopted from O’Donnell et al. (2010). PCR products were purified using an ExoSap purification protocol. Purified amplification products were sequenced by the Macrogen sequencing facilities (Macrogen, Seoul, South Korea).

### Phylogenetic analyses

Raw sequences were assembled using Geneious Prime (Biomatters, New Zealand). Automatic alignment was conducted with MAFFT (Katoh et al. 2002) using an E-INS-i algorithm, a 100PAM/k=2 scoring matrix, a gap open penalty of 1.3 and an offset value of 0.123. Manual fine-tuning of the aligned dataset was performed in Geneious Prime.

*Fusarium* sequence data of *rpb1*, *rpb2* and *tef1a* was extracted from GenBank (September 20, 2020) using the ‘NCBI Nucleotide extraction’ tool in Geneious Prime. Together with the newly generated sequences for the 19 Vietnamese *Fusarium* wilt accessions, the total sequence data matrix consisted of 542 specimens divided over 210 species (Suppl. Table S1). Of those, 11 species belonging to different closely related genera of *Fusarium* within the Nectriaceae family were chosen as outgroup (*Cosmospora, Cylindrocarpon, Fusicolla, Macroconia* and *Microcera*). Newly generated sequences were deposited in the GenBank sequence database (Table 1). Furthermore, in order to compare the newly collected Vietnamese *Foc* accessions with the known Vegetative Compatibility Groups (VGC’s), the sequence dataset included *Foc* samples representing all VCG’s (see Table S1; Ordonez et al. 2015), except for VCG01212 and VCG0129. For the latter two, only one locus was available causing phylogenetic biases due to the occurrence of too much missing data.

Possible incongruency between the different datasets was inferred by conducting an ILD test (Farris et al. 1995) as implemented in PAUP* v.4.0b10 (Swofford 2003) with following parameters applied: simple taxon addition, TBR branch swapping and heuristic searches of 1000 repartitions of the data. Despite the well-known sensitivity of the ILD test (Barker and Lutzoni 2002), the results of this test were compared in light of the resolution and support values for each of the single gene topologies. As a result, possible conflict between data matrices was visually inspected by searching for conflicting relationships within each topology (obtained per single sequence data matrix) that were supported by a Maximum Likelihood (ML) support value >70% (hard vs. soft incongruence; Johnson and Soltis 1998; Pirie 2015). A conflict was assumed to be significant if two different relationships for the same set of taxa (one being monophyletic and the other non-monophyletic) were observed in rival trees. ML analyses were conducted under the RAxML search algorithm (Stamatakis 2014) with the GTRGAMMAI approximation of rate heterogeneity for each gene. ML bootstrapping was carried out on five hundred bootstrapped datasets using the RAxML Rapid bootstrap algorithm (ML-BS).

The best-fit nucleotide substitution model for each dataset was selected using jModelTest 2.1.4. (Posada 2008) out of 88 possible models under the Akaike information criterion (AIC). The GTR+I+G model was determined as best fit for *rpb1*, while the TVM+G model was calculated as best substitution model for *tef1a* and HKY+I+G as best substitution model for *rpb2*. Consequently, we used a mixed-model approach to apply different evolutionary models on each DNA region of the combined dataset (Ronquist and Huelsenbeck 2003). Bayesian inference analyses were conducted with MrBayes v3.2.6 (Ronquist et al. 2012) on three individual data partitions and a combined data matrix. Each analysis was run two times for 20 million generations. Trees were sampled every 5000^th^ generation. Chain convergence and ESS parameters were inspected with TRACER v.1.4 (Rambaut and Drummond 2007). Only nodes with Bayesian posterior probabilities (BPP) above 0.95 were considered as well supported by the data (Suzuki et al. 2002).

## Results

Pathogenic *Fusarium* wilt infections are prevalent in most of northern Viet Nam as they have been observed in all provinces of northern Viet Nam that were sampled in this study. The 19 *Fusarium* wilt infections collected based on the typical plant Fusariosis symptoms (old leaves turning yellow, leaves gradually collapsing, petioles broken close to the midrib with dead leaves remaining attached to the pseudostem, pseudostem sheaths longitudinally splitting near the base, vascular necrosis) were cultured and further morphologically and molecularly analysed.

Morphological characterization of the cultured pathogenic *Fusarium* wilt isolates showed that when the isolates were grown on CLA medium, they produced macroconidia that were uniform in size and form. On SNA medium, the morphology of the macroconidia was sometimes less uniform in size compared to when SLA medium was used. Except for two accessions (FOC56 and FOC61), no aroma was observed among the pathogenic *Fusarium* isolates collected in northern Viet Nam. In general, for all isolates, we observed that macroconidia are sickle-shaped, 3-7 septate, and thin-walled. Microconidia are oval to kidney-shaped, 0-1 septate. Chlamydospores were round and thick-walled. Subtle differences have been observed in the colony morphology and coloration. Based on these morphological differences, we tried to identify different groups within the *Fusarium* isolates analysed. The first group, consisting of 14 isolates (FOC1, 2, 5, 6-1, 7, 11, 16, 18, 21, 23-2, 24, 25-1, 25-2, 38), is characterized by a purple reverse in the centre, white-greyish towards the periphery. The colony surface is dry and is filamentous at the edge. On CLA medium, it produces ample macroconidia, yet only little microconidia. On PDA and SNA medium, it produces prolific microconidia. The second group has a reverse colony colour containing a small touch of dark purple in the centre, gradually discolouring to white towards the edge. This type is observed for isolates FOC 4 and 10. The surface of these colonies is also dry and filamentous at the margin. On CLA medium, ample macroconidia are produced whereas on PDA and SNA medium, the presence of macroconidia is less profound. On the latter two media, prolific microconidia are produced. A third group of isolates (FOC56 and 61) is characterized by an unpigmented, white colony reverse and a dry colony surface with filamentous margin. On CLA medium, a large number of macroconidia is produced while on PDA and SNA medium macroconidia are hardly formed. On PDA and SNA prolific microconidia are produced, whereas on CLA medium only few microconidia were observed. In addition, FOC 56 and 61 isolates are characterized by a typical strong odour of the older cultures. FOC 58 falls a bit amidst the first and second group, containing a pale purple colony reverse colour that becomes whitish towards the periphery and with a dry colony surface appearance.

### Phylogenetic analyses of pathogenic *Fusarium* wilt isolates

Sequence characteristics of all data matrices analysed are summarized in Table 2. Despite the fact that sometimes not all gene markers could be sequenced, their absence did not influence the overall phylogenetic results, as sufficient nucleotide variation was present. No significant incongruence between all three sequence datasets (with all *P*-value being larger than 0.05) was found using the partition homogeneity test. Visual examination of the two different partitions of the combined dataset corroborates this congruency analysis.

**Table 2.** Alignment and sequence characteristics of the different partitions (including outgroup specimens).

|  | *rpb1* | *rpb2* | *tefla* |
| --- | --- | --- | --- |
| N° taxa | 469 | 539 | 273 |
| Sequence length range | 558-1574 | 597-859 | 343-636 |
| Aligned sequence range | 1578 | 917 | 797 |
| Variable characters | 1106 (70%) | 573 (62%) | 529 (66%) |
| Constant characters | 472 | 344 | 268 |

Phylogenetic analyses of the 19 pathogenic *Fusarium* wilt isolates that were found on various northern Vietnamese bananas showed that although overall morphological characterization pointed towards *Fusarium oxysporum* f. sp. *cubense*, it was clear that they had various evolutionary origins (Fig. 3c, d). Of the 19 accessions analysed, two (FOC61 and FOC56) were placed within the *Fusarium odoratissimum* clade (as defined by Maryani et al. 2019a, Fig. 3d) and for which pathogenicity tests by Maryani et al. (2019a) showed that the members of this group caused infections in Cavendish and Gros Michel AAA banana varieties. In addition, VCG 01213 and VCG 01216 are positioned close to FOC61 and FOC56. As a result, two of the 19 (10.5%) northern Vietnamese pathogenic *Fusarium* wilt isolates are assumed to be *Foc-*TR4 (also taking the morphological characterization into account). Interestingly, one of the two isolates characterized as *Foc-*TR4 (FOC61) infected a Cavendish plantation in Vinh Phuc province, whereas the other infection of *Foc-*TR4 (FOC56) took place on ABB Tay banana cultivars situated on a smallholder farm in Nam Dinh province (Table 1). The largest group (13 accessions; 68.5%) of pathogenic *Fusarium* wilt isolates in northern Vietnamese bananas belong to the recently delineated *Fusarium tardichlamydosporum* clade (Fig. 3d). Pathogenicity tests carried out for this clade by Maryani et al. (2019a) have indicated a large infection rate in Gros Michel cultivars for this lineage and therefore members of the *Fusarium tardichlamydosporum* clade are consequently classified as *Foc*-Race 1. Furthermore, the isolates that fell within the *Fusarium tardichlamydosporum* were also most closely related to VCG 0125, a known *Foc*-Race 1 representative. In northern Viet Nam, infections of *Foc*-Race 1 occurred both on wild and cultivated accessions. For the cultivated accessions, the *Foc*-Race 1 was only found on the ABB Tay banana cultivar, yet was clearly spread in northern Viet Nam as it was found in eight different provinces (Ha Giang, Yen Bai, Lao Cai, Bac Giang, Nam Dinh, Ha Nam, Tuyen Quang, Ha Noi; Table 1). Most interestingly, *Foc*-Race 1 was also identified (isolate FOC5) in an individual of the wild banana *Musa lutea* (section Callimusa). Here the infected accession grew sympatrically with other individuals of *Musa lutea* as well as with *Musa itinerans*. The area where this infection occurred was a steep, abandoned rice terrace in Yen Bai province where hundreds of individuals of both wild species co-occurred and was rather close to one of the smallholder farms where *Foc*-Race 1 was also detected (isolate FOC6-1). In addition to the *Foc*-Race 1 infections caused by *Fusarium tardichlamydosporum*, also *F. duoseptatum* is classified as a *Foc*-Race 1 *Fusarium* wilt (see Maryani et al. 2019). An infection of this latter *Foc* isolate (FOC38) was found only once in northern Viet Nam (Nghe An province; c. 5% of the *Fusarium* wilt infections) where it infected the ABB Tay banana cultivars that where grown on a smallholder farm. The VCG’s that occurred in the same clade as FOC38 are VCG 01223 and VCG 01217, with the latter being known as a *Foc*-Race 1 representative (e.g. Katan 1999, Fraser-Smith et al. 2014).

**Fig. 3.**
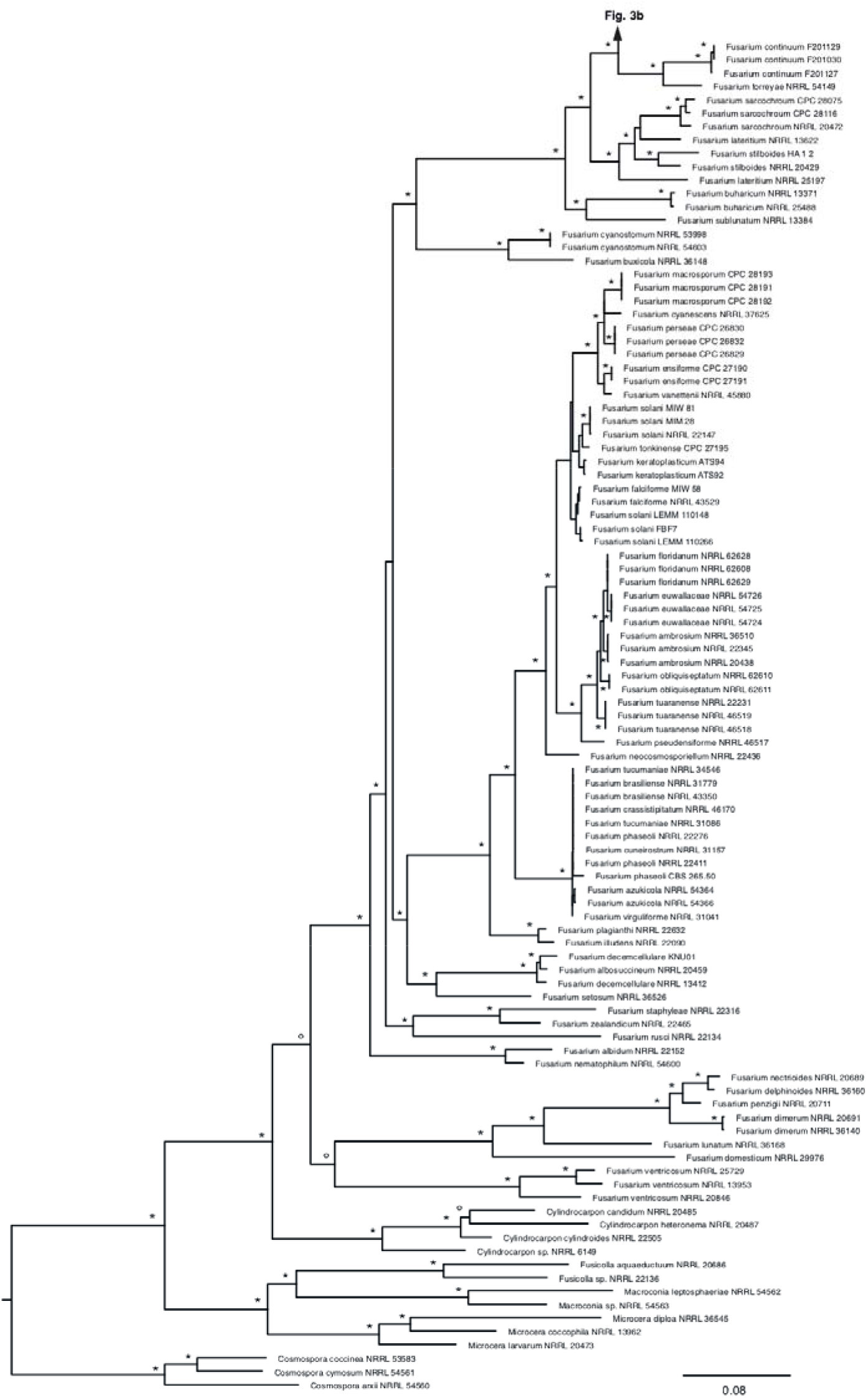
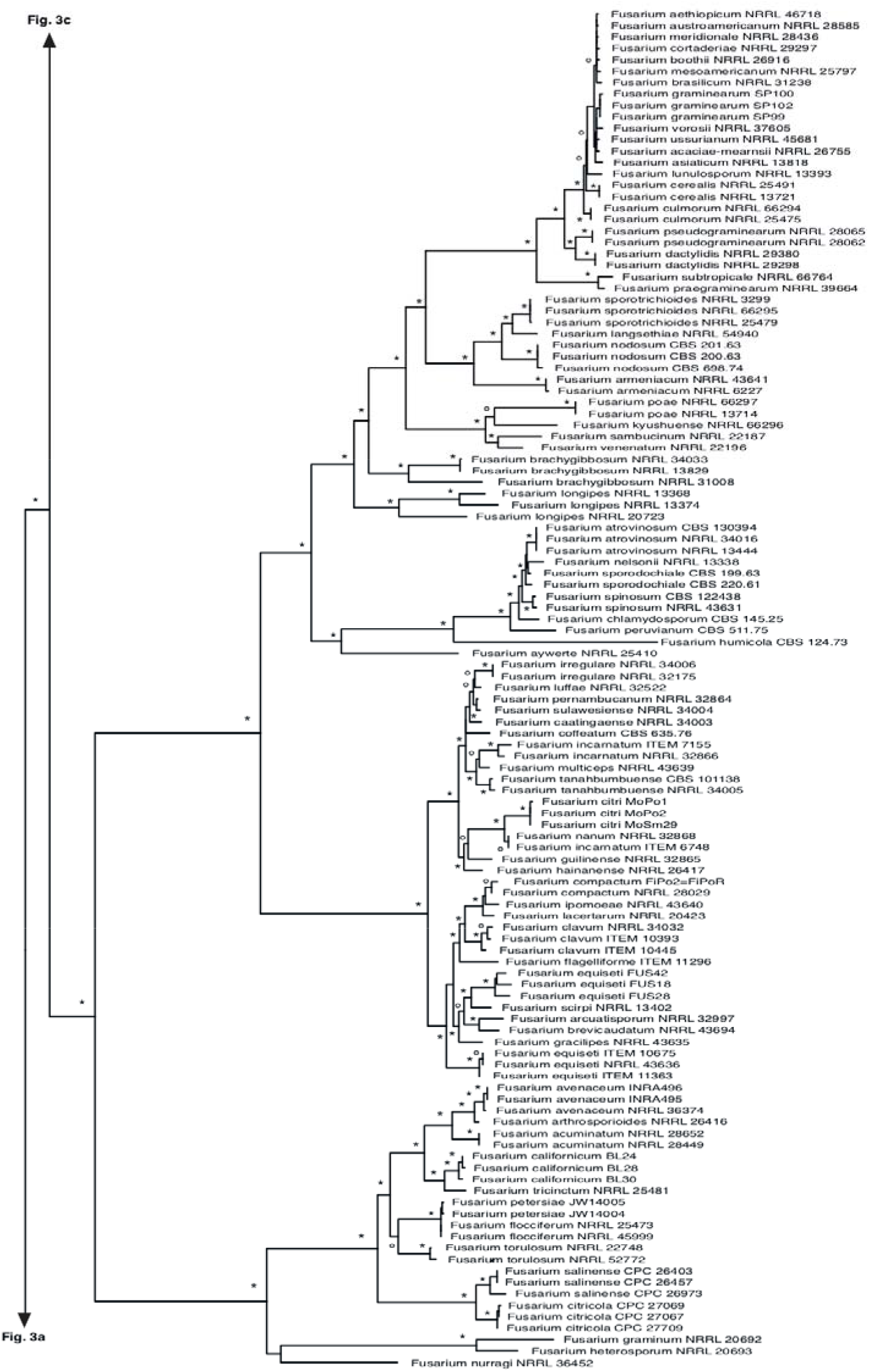
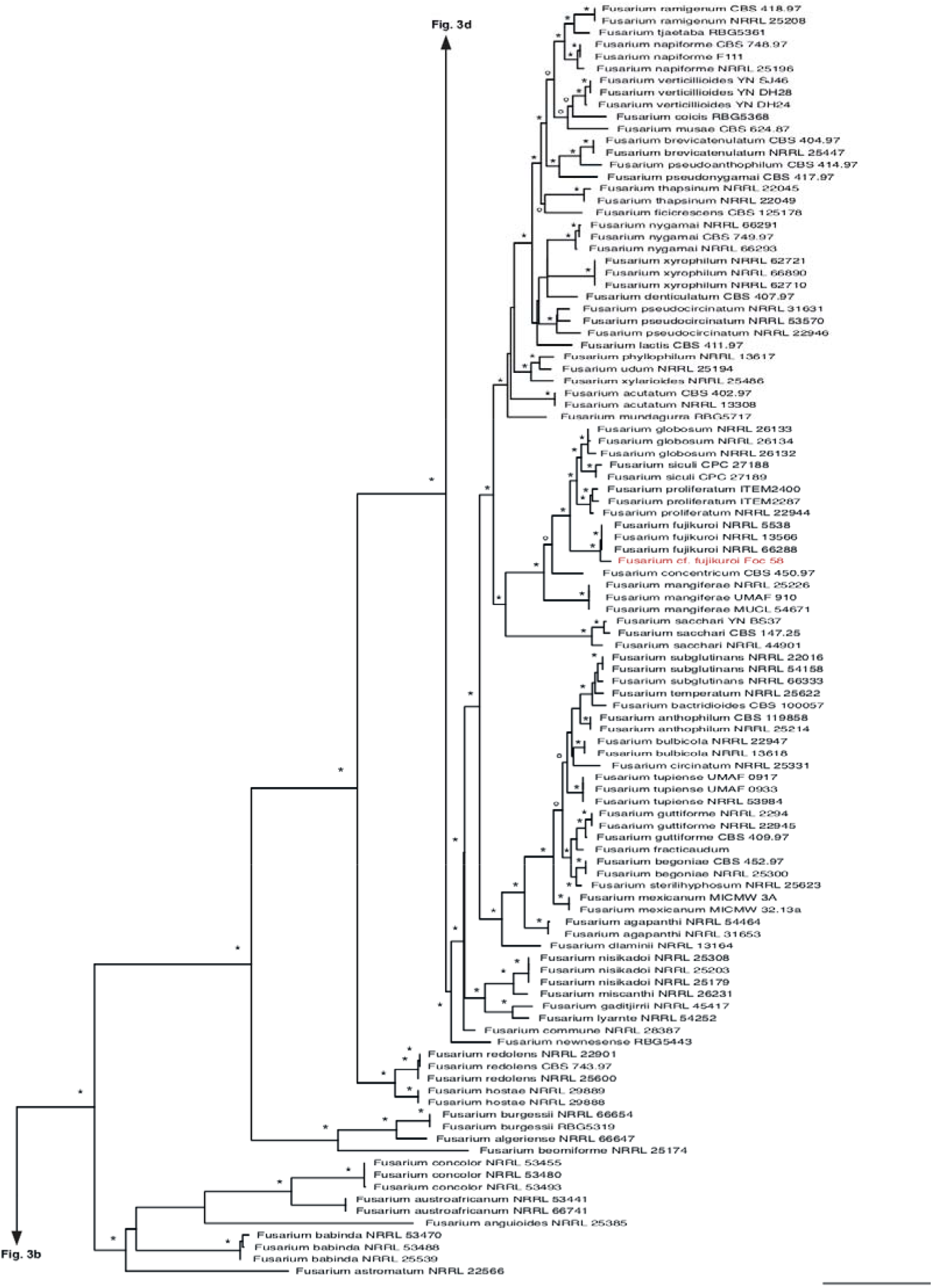
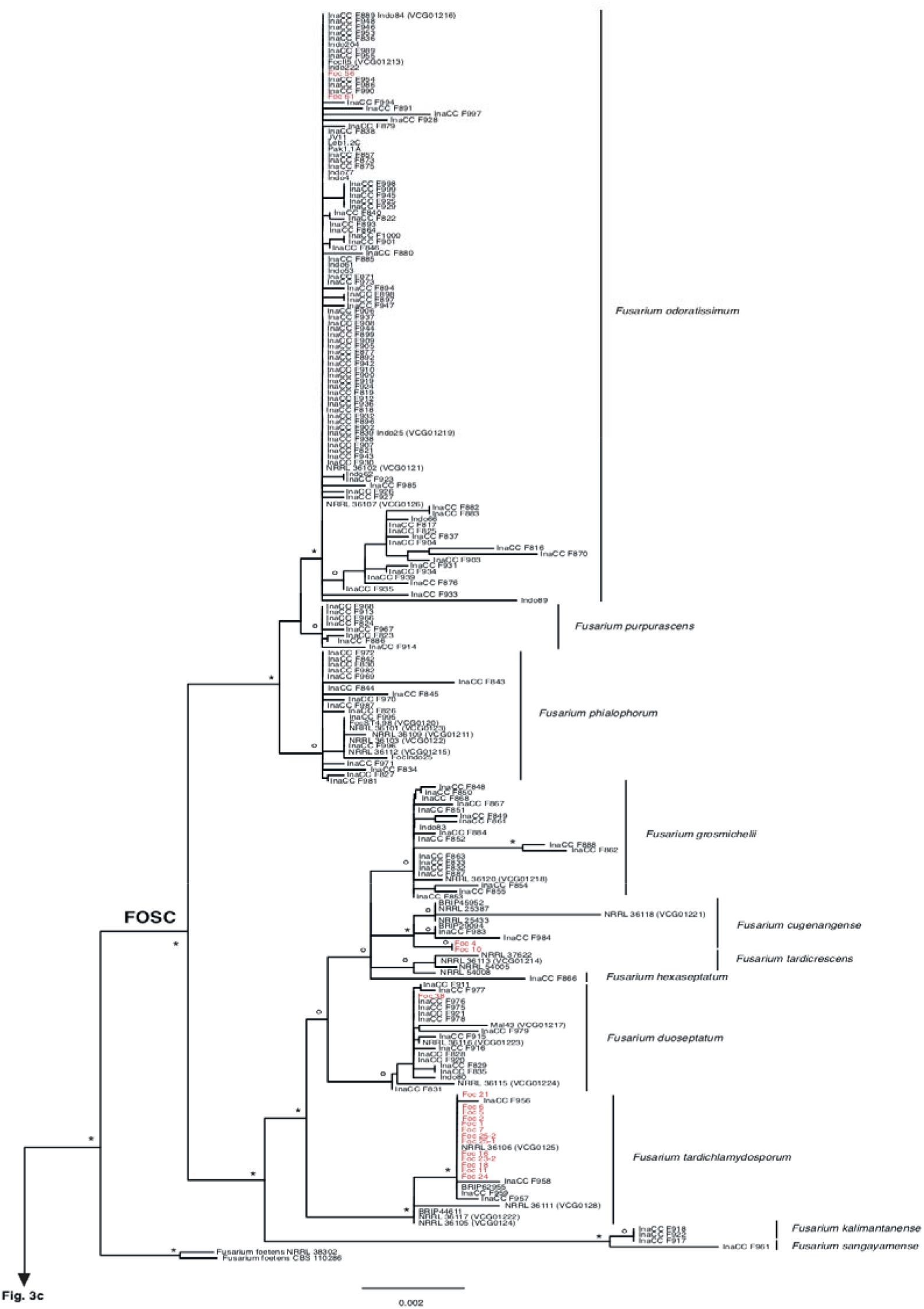
Maximum Likelihood topology obtained via heuristic search algorithm of the combined *rpb1*, *rpb2* and *tef1a* data matrix. Bootstrap support (ML-BS) values above 50 are indicated with a dot, ML-BS values above 75 are indicated with an asterisk. No indication above the branches indicates a ML-BS value below 50. Newly included accessions are indicated in red. FOSC: *Fusarium oxysporum* species complex, FFSC: *Fusarium fujikuroi* species complex.

In addition to the *Foc*-Race 1 and *Foc-*TR4 infections, two pathogenic *Foc* isolates (FOC4 and FOC10) were found in northern Viet Nam (10.5%) that belong to the recently described *F. cugenangense* (Maryani et al. 2019a; Fig. 3d). Up to now, this *Fusarium* species was considered to be strictly Indonesian (see Maryani et al. 2019a). Pathogenicity tests conducted for representatives of *F. cugenangense* by Maryani et al. (2019a) have demonstrated that it only causes a mild infection in Gros Michel and Cavendish and were regarded as non-pathogenic for the above-mentioned AAA cultivars. However, our results clearly show that the infection of this isolate also occurred on ABB Tay banana cultivars in northern Viet Nam, where it had a large impact on the fitness of the infected host plants. Although additional confirmation is needed, Maryani et al (2019a) assume that representatives of the *F. cugenangense* clade should be considered as *Foc*-Race 2 (Maryani et al. 2019a; Fig. 3d), yet more thorough analyses need to be carried out in order to further confirm this hypothesis. The VCG that occurred in the same clade as FOC4 and FOC10 is VCG 01221.

A final pathogenic *Fusarium* wilt infection (FOC58) that was regarded upon collection in the field and during morphological screening as a *Foc* infection (c. 5% of the *Fusarium* wilt infections observed in this study), was in fact not situated in the *Fusarium oxysporum species complex* (FOSC) but was a distinct lineage sister to *Fusarium fujikuroi* (Fig. 3c), a well-known pathogen of rice (e.g. Wulff et al. 2010; Choi et al. 2018). This pathogenic *Fusarium* isolate was the prime infection source of ABB Tay bananas that were cultivated on a small plantation for local use in Nam Dinh province (Fig. 1).

## Discussion

### Fusarium wilting in Vietnam: lineage identification

To better manage the significant threat of *Foc* dispersion in the northern Viet Nam, the correct identification and abundancy of the *Foc* strains that cause Fusarium wilting in bananas in the region is necessary. This is the basis for eradication-confinement and suppression-contention measures (Perez-Vicente et al. 2014). Since the survey of Vakili et al. (1968), *Foc* Race 1 has been considered as the main *Foc* infecting edible bananas in Viet Nam. With the emergence of *Foc* TR4 it remained unclear how abundant this new pathogenic *Foc* strain had become in Viet Nam. Although officially present in Viet Nam for a few years (Hung et al. 2017; Zheng et al. 2018), *Foc* TR4 was already observed on Cavendish bananas in 1998 in Southern China (Hu et al. 2006). A few years later, in 2002, *Foc* TR4 was also found in Chinese regions adjacent to northern Viet Nam (Hu et al. 2006; Li et al. 2013). With the current shift of Cavendish cultivation in Asia from China to its neighbouring countries Laos, Myanmar and Viet Nam, there is also an active spread of *Fusarium* pathogens through transportation of planting material, farming equipment and contaminated soil from China (Zheng et al. 2018), so that *Foc* TR4 can quickly become the most dominant *Foc* race in Viet Nam affecting banana cultivation.

The present study applies the FOSC species delimitation concept of Maryani et al. (2019) in order to more thoroughly delineate the *Foc* lineages that were sampled in northern Viet Nam. Furthermore, the incorporation of the different VCG’s in the current phylogenetic dataset allowed us to link the Vietnamese *Foc* isolates with the one of the currently known VCG’s that have been assessed in the past. Based on the compatibility of the novel material with the VCG’s that are present in the same clade, as well as with ther specific species allocation following the species delineation concept of Maryani et al. (2019), we classified the northern Vietnamese *Foc* isolates into one of the known *Foc* Races. Accordingly, our results shows that *Foc* Race 1 is the most common isolate in northern Viet Nam, causing 74% of all the infections. A more in-depth molecular characterization shows that among these *Foc* Race 1 infections, 13 out of 14 isolates are caused by the species *Fusarium tardichlamydosporum* and one by the species *F. duoseptatum*. Interestingly, *Foc* Race 1 *Fusarium* wilting caused by *F. tardichlamydosporum* occurred in different provinces than *Foc* Race 1 *Fusarium* wilting caused by *F. duoseptatum* (Fig. 1). Whereas *F. tardichlamydosporum* is commonly present throughout the northernly oriented Northeastern region and Red River Delta, *F. duoseptatum* is not present in these more northerly oriented geographic regions but occurs in the more centrally oriented North Central region in Viet Nam. Also, from a global distributional perspective, *F. tardichlamydosporum* is much more widespread than *F. duoseptatum*, with the first species located in Australia, Indonesia, Malaysia, Honduras and Brazil, and the latter only known to date from Indonesia and Malaysia (Maryani et al. 2019a). Compared to *Foc*-Race 1, *Foc-*TR4 infections (*F. odoratissimum*; Maryani et al. 2019a) account for only 10% of the Fusarium wilting in northern Viet Nam demonstrating that *Foc-TR4* has not yet become a dominant banana pathogen, unlike other countries in Asia where there is a tendency to grow Cavendish cultivars as large monocultures, such as in China, the Philippines and Taiwan. The *Foc-*TR4 isolates that were found in the current study were located in the River Delta region of northern Vietnam, provinces Vinh Phuc and Nam Dinh, which are rather distant from *Foc-*TR4 infected regions in Southern China. This indicates a gradual spreading in Viet Nam of *F. odoratissimum* (=*F. oxysporum* f. sp. *cubense* TR4) towards the south as the TR4 isolates analysed by Zheng et al. (2018) were collected in the upper North of Viet Nam in the Lao Cai province at only few kilometres from the border with China (Yunnan). At the moment it is unclear whether the occurrence of *Foc-*TR4 in Viet Nam is still in an initial lag phase, with the potential of largely increasing its distribution range in the country if conditions would improve for the disease to spread (Pegg et al. 2019). Especially the replacement of citrus plantations and maize fields by Cavendish monocultures provides an ideal basis for *Foc-*TR4 to rapidly spread as plants available for infection become less limited. A more worrying observation is that *Foc-*TR4 is not only found in Cavendish bananas in Viet Nam, but that it also poses a threat to local banana varieties as is observed in the current study where ABB Tay banana cultivars seem to be prone to *Fusarium* wilting caused by *Foc-*TR4.

The current study demonstrates that *Foc*-Race 2 infections of Vietnamese bananas also account for 10% of the *Fusarium* wilting in northern Viet Nam. In general, *Foc*-Race 2 infections occur on triploid ABB Bluggoe varieties and its closely related cooking cultivars (Jones, 2000). Besides Bluggoe cooking bananas, *Foc*-Race 2 also infects the tetraploid AAAA Bodles Altafort hybrid between Gros Michel (AAA) and Pisang Lilin (AA)(Stover and Simmonds 1987). In addition, experimental infection of *Ensete ventricosum* demonstrated that this important Ethiopian crop is highly susceptible to *Foc*-Race 2 (Ploetz, 2005). With the confirmation of a *Foc*-Race 2 infection affecting also representatives of the ABB Tay banana cultivar, it is clear that *Fusarium* wilting caused by *Foc*-Race 2 is potentially more widespread than has often been assumed.

### Fusarium cf. fujikuroi as a novel pathogen of bananas

In addition to the pathogenic *Fusarium* isolates collected from northern Vietnamese bananas belonging to the FOSC, an infection with symptoms similar to *Foc* wilting was observed, yet the cultured isolate did not belong to FOSC. The morphological colony characteristics were very comparable to those observed for FOSC cultures by having a pale purple colony reverse colour that became whitish towards the periphery with a dry colony surface appearance. However, when genetically assessing within the *Fusarium* genus, this isolate did not fall within *F. oxysporum* representatives, but was member of the *Fusarium fujikuroi* species complex (FFSC) where it is the sister lineage of *F. fujikuroi*. It is not uncommon that several *Fusarium* species cause the same disease pattern as this phenomenon has also been identified in mango deformity (Lima et al. 2009) and sugar beet wilting (Burlakoti et al. 2012). Within the FFSC, some species are known to be pathogenic for some *Musa* cultivars (*F. proliferatum, F. verticillioides, F. sacchari, F. lumajangense, F. desaboruense* and *F. musae*; Maldonado-Bonilla et al. 2019, Van Hove et al. 2002, Huang et al. 2019, Maryani et al. 2019b), yet to date no other species of the *Fusarium fujikuroi* species complex - except for the abovementioned -was identified as pathogen for *Musa*. From a phylogenetic point of view, the novel pathogenic *Fusarium* isolate that infected a triploid ABB Tay banana cultivar in Nam Dinh province is sister to *F. fujikuroi. Fusarium fujikuroi* is a widespread phytopathogen causing the bakanae disease in various *Oryza sativa* cultivars (rice), but is also known to have a major impact on many other economically important crops (e.g., maize, wheat). However, to date, no infection of *F. fujikuroi* or a close relative has been detected in bananas. This result increases our knowledge on the diversity of *Fusarium* species that cause wilting symptoms on bananas. More importantly, it also demonstrates the urgent need for an accurate identification of plant pathogens that are morphologically very difficult to distinguish from each other in the field.

### Fusarium wilting on wild bananas

Although mainly observed on cultivated bananas, *Foc* has also been rarely recorded on wild Musa species (Ploetz and Pegg 2000). Waite (1954) noticed that *Fusarium* wilting also occurred on *M. acuminata, M. balbisiana, M. schizocarpa* and *M. textilis*. Since these specific *Musa* species belong to different sections in the genus - section *Musa* and *Callimusa* (*Australimusa*) - it is therefore likely that *Foc* can also infect other wild bananas. The current finding of a *Foc*-Race 1 infection on a wild representative of *Musa lutea*, could indicate that wild species are perhaps more susceptible to *Fusarium* wilting than previously assumed. Besides this one individual showing visual symptoms of *Fusarium* wilting, none of the hundreds of individuals of *Musa itinerans* and *Musa lutea* surveyed in the same population showed any sign of *Foc* infection. This lack of visual symptoms either implies that *F. oxysporum* f. sp*. cubense* could have been absent from all those other wild accessions or that the *Foc*-Race 1 pathogen was present but failing to cause the disease in the other wild bananas. If the latter assumption is true, this could indicate that the pathogen not necessarily co-evolved together with its host in Southeast Asia as postulated by Vakili, (1965) but that *Fusarium oxysporum* is omnipresent throughout the native distribution range of the *Musa* genus and that infections take place when the plant is weakened due to external biotic of abiotic stressors and the endophytic equilibrium is disturbed.

## Acknowledgements

This study was funded by a bilateral grant between the Research Foundation - Flanders (FWO) and the Vietnamese National Foundation for Science and Technology Development (NAFOSTED) G0D9318N / FWO.106-NN.2017.02. This work was also supported by the University of Queensland via the Bill & Melinda Gates Foundation project ‘BBTV mitigation: Community management in Nigeria, and screening wild banana progenitors for resistance’ [OPP1130226]. We are grateful to for the technical laboratory work carried out at Meise Botanic Garden by Wim Baert, Pieter Asselman, Lynn Delgat and Annelies Heylen. The authors thank all donors who supported this work also through their contributions to the CGIAR Fund (http://www.cgiar.org/who-we-are/cgiar-fund/fund-donors-2/), and in particular to the CGIAR Research Program Roots, Tubers and Bananas (RTB-CRP).

## Author Contribution

LLT, AM, DTV and SBJ conceptualized the manuscript. LLT, SdB, AM and SBJ carried out the experimental work. SBJ, AM and LLT wrote the original manuscript. MA, SG and CD optimized the morphological analyses. RS, FV, DTV and SBJ acquired funding for the project. All other authors optimized the initial draft and provided helpful contributions to the finalization of the paper.

## Supplementary Tables

**Supplementary Table S1.** List of accessions used for the phylogenetic analyses, including voucher information and GenBank numbers. Asterisks indicate accessions for which new sequences were generated in the current study.

| *Species* | Accession n° | *rpb1* | *rpb2* | *tefla* |
| --- | --- | --- | --- | --- |
| *Cosmospora arxii* | NRRL 54560 | JX171554 | JX171666 | - |
| *Cosmospora coccinea* | NRRL 53583 | JX171545 | JX171657 | - |
| *Cosmospora cymosum* | NRRL 54561 | JX171555 | JX171667 | - |
| *Cylindrocarpon candidum* | NRRL 20485 | JX171588 | JX171474 | - |
| *Cylindrocarpon cylindroides* | NRRL 22505 | JX171499 | JX171612 | - |
| *Cylindrocarpon heteronema* | NRRL 20487 | JX171475 | JX171589 | - |
| *Cylindrocarpon sp.* | NRRL 6149 | JX171445 | JX171559 | - |
| *Fusarium acaciae/mearnsii* | NRRL 26755 | KM361640 | KM361658 | - |
| *Fusarium acuminatum* | NRRL 28449 | MG282373 | MG282402 | - |
| *Fusarium acuminatum* | NRRL 28652 | MG282384 | MG282414 | - |
| *Fusarium acutatum* | CBS 402.97 | MT010947 | KT154005 | MT010989 |
| *Fusarium acutatum* | NRRL 13308 | MN193911 | MN193883 | - |
| *Fusarium aethiopicum* | NRRL 46718 | KM361652 | KM361670 | - |
| *Fusarium agapanthi* | NRRL 31653 | KU900619 | KU900624 | - |
| *Fusarium agapanthi* | NRRL 54464 | KU900622 | KU900627 | - |
| *Fusarium albidum* | NRRL 22152 | JX171492 | JX171605 | - |
| *Fusarium albosuccineum* | NRRL 20459 | JX171585 | JX171585 | - |
| *Fusarium algeriense* | NRRL 66647 | MF120488 | MT409451 | - |
| *Fusarium ambrosium* | NRRL 20438 | JX171470 | JX171584 | - |
| *Fusarium ambrosium* | NRRL 22345 | KC691586 | KC691618 | - |
| *Fusarium ambrosium* | NRRL 36510 | KC691588 | KC691619 | - |
| *Fusarium anguioides* | NRRL 25385 | JX171624 | JX171624 | - |
| *Fusarium anthophilum* | CBS 119858 | MT010940 | KU604275 | MT010997 |
| *Fusarium anthophilum* | NRRL 25214 | KU171696 | KF466403 | - |
| *Fusarium arcuatissporum* | NRRL 32997 | HM347164 | GQ505802 | - |
| *Fusarium armeniacum* | NRRL 43641 | HM347192 | GQ505494 | - |
| *Fusarium armeniacum* | NRRL 6227 | JX171446 | HQ154480 | - |
| *Fusarium arthrosporioides* | NRRL 26416 | MG282383 | MG282413 | - |
| *Fusarium asiaticum* | NRRL 13818 | JX171459 | JX171573 | - |
| *Fusarium astromatum* | NRRL 22566 | JX171500 | JX171613 | - |
| *Fusarium atrovinosum* | CBS 130394 | MN120714 | MN120734 | - |
| *Fusarium atrovinosum* | NRRL 13444 | JX171454 | JX171568 | - |
| *Fusarium atrovinosum* | NRRL 34016 | HM347170 | GQ505475 | - |
| *Fusarium austroafricanum* | NRRL 53441 | MH742536 | MH742615 | - |
| *Fusarium austroafricanum* | NRRL 66741 | MH742537 | MH742616 | - |
| *Fusarium austroamericanum* | NRRL 28585 | KM361643 | KM361661 | - |
| *Fusarium avenaceum* | INRA495 | MH667523 | MH667549 | - |
| *Fusarium avenaceum* | INRA496 | MH667524 | MH667550 | - |
| *Fusarium avenaceum* | NRRL 36374 | MG282366 | MG282395 | - |
| *Fusarium aywerte* | NRRL 25410 | JX171513 | JX171626 | - |
| *Fusarium azukicola* | NRRL 54364 | KJ511276 | KJ511287 | - |
| *Fusarium azukicola* | NRRL 54366 | KJ511277 | KJ511288 | - |
| *Fusarium babinda* | NRRL 25539 | JX171632 | JX171632 | - |
| *Fusarium babinda* | NRRL 53470 | MH742548 | MH742627 | - |
| *Fusarium babinda* | NRRL 53488 | MH742552 | MH742631 | - |
| *Fusarium bactridioides* | CBS 100057 | MT010939 | MT010963 | MT010995 |
| *Fusarium begoniae* | CBS 452.97 | MT010936 | MT010964 | MT010998 |
| *Fusarium begoniae* | NRRL 25300 | MN193914 | MN193886 | - |
| *Fusarium beomiforme* | NRRL 25174 | JX171506 | JX171619 | - |
| *Fusarium boothii* | NRRL 26916 | KM361641 | GQ915487 | - |
| *Fusarium brachygibbosum* | NRRL 13829 | JX171460 | JX171574 | - |
| *Fusarium brachygibbosum* | NRRL 31008 | JX171529 | MH845433 | - |
| *Fusarium brachygibbosum* | NRRL 34033 | HM347172 | GQ505482 | - |
| *Fusarium brasiliicum* | NRRL 31238 | KM361645 | KM361663 | - |
| *Fusarium brasiliense* | NRRL 31779 | KJ511272 | KJ511283 | - |
| *Fusarium brasiliense* | NRRL 43350 | KJ511274 | KJ511285 | - |
| *Fusarium brevicatenulatum* | CBS 404.97 | MT010948 | MT010979 | MT011005 |
| *Fusarium brevicatenulatum* | NRRL 25447 | MN193915 | MN193887 | - |
| *Fusarium brevicaudatum* | NRRL 43694 | HM347193 | GQ505846 | - |
| *Fusarium buharicum* | NRRL 13371 | JX171449 | JX171563 | - |
| *Fusarium buharicum* | NRRL 25488 | KX302920 | KX302928 | - |
| *Fusarium bulbicola* | NRRL 13618 | KF466394 | KF466404 | - |
| *Fusarium bulbicola* | NRRL 22947 | KU171679 | KU171699 | - |
| *Fusarium burgessii* | NRRL 66654 | MF120495 | MT409450 | - |
| *Fusarium burgessii* | RBG5319 | KJ716217 | HQ646392 | - |
| *Fusarium buxicola* | NRRL 36148 | JX171534 | HM068357 | - |
| *Fusarium caatingaense* | NRRL 34003 | HM347166 | GQ505805 | - |
| *Fusarium californicum* | BL24 | MK878580 | MK878565 | - |
| *Fusarium californicum* | BL28 | MK878582 | MK878567 | - |
| *Fusarium californicum* | BL30 | MK878584 | MK878569 | - |
| *Fusarium cerealis* | NRRL 13721 | KM361638 | KM361656 | - |
| *Fusarium cerealis* | NRRL 25491 | MG282371 | MG282400 | - |
| *Fusarium cf. fujikuroi* * | Foc 58 | XX000000 | XX000000 | XX000000 |
| *Fusarium chlamydosporum* | CBS 145.25 | MN120715 | MN120735 | - |
| *Fusarium circinatum* | NRRL 25331 | JX171510 | HM068354 | - |
| *Fusarium citri* | MoPo1 | - | LT970750 | LT970778 |
| *Fusarium citri* | MoPo2 | - | LT970751 | LT970779 |
| *Fusarium citri* | MoSm29 | - | LT970754 | LT970782 |
| *Fusarium citricola* | CPC 27067 | LT746287 | LT746307 | LT746194 |
| *Fusarium citricola* | CPC 27069 | LT746288 | LT746308 | LT746195 |
| *Fusarium citricola* | CPC 27709 | LT746289 | LT746309 | LT746196 |
| *Fusarium clavum* | ITEM 10393 | - | LN901601 | LN901566 |
| *Fusarium clavum* | ITEM 10445 | - | LN901603 | LN901568 |
| *Fusarium clavum* | NRRL 34032 | HM347171 | GQ505813 | - |
| *Fusarium coffeatum* | CBS 635.76 | MN120717 | KU604328 | - |
| *Fusarium coicis* | RBG5368 | KP083269 | KP083274 | - |
| *Fusarium commune* | NRRL 28387 | JX171638 | HM068356 | - |
| *Fusarium compactum* | FiPo2=FiPoR | - | LT970748 | LT970776 |
| *Fusarium compactum* | NRRL 28029 | HM347150 | GQ505780 | - |
| *Fusarium concentricum* | CBS 450.97 | MT010942 | MT010981 | MT010992 |
| *Fusarium concolor* | NRRL 53455 | MH742506 | MH742583 | - |
| *Fusarium concolor* | NRRL 53480 | MH742513 | MH742591 | - |
| *Fusarium concolor* | NRRL 53493 | MH742535 | MH742614 | - |
| *Fusarium continuum* | F201030 | KM520387 | KM236782 | - |
| *Fusarium continuum* | F201127 | KM520386 | KM236779 | - |
| *Fusarium continuum* | F201129 | KM520385 | KM236781 | - |
| *Fusarium cortaderiae* | NRRL 29297 | KM361644 | KM361662 | - |
| *Fusarium crassistipitatum* | NRRL 46170 | KJ511275 | KJ511286 | - |
| *Fusarium cugenangense* | InaCC F983 | LS479559 | LS479307 | LS479756 |
| *Fusarium cugenangense* | InaCC F984 | LS479560 | LS479308 | LS479757 |
| *Fusarium cugenangense* | NRRL 25433 | LS479462 | LS479202 | LS479648 |
| *Fusarium cugenangense* | NRRL 36118 (VCG01221) | LS479477 | LS479221 | LS479669 |
| *Fusarium cugenangense* | BRIP29094 | KX434922 | KX434957 | - |
| *Fusarium cugenangense* | BRIP45952 | KX434923 | KX434958 | - |
| *Fusarium cugenangense* | NRRL 25387 | JX171625 | HM347209 | - |
| *Fusarium cugenangense* * | Foc 10 | - | XX000000 | XX000000 |
| *Fusarium cugenangense* * | Foc 4 | - | XX000000 | XX000000 |
| *Fusarium culmorum* | NRRL 25475 | JX171515 | JX171628 | - |
| *Fusarium culmorum* | NRRL 66294 | MG282380 | MG282410 | - |
| *Fusarium cuneirostrum* | NRRL 31157 | KJ511271 | FJ240389 | - |
| *Fusarium cyanescens* | NRRL 37625 | HM347175 | - | - |
| *Fusarium cyanostomum* | NRRL 53998 | JX171546 | JX171658 | - |
| *Fusarium cyanostomum* | NRRL 54603 | JX171553 | JX171665 | - |
| *Fusarium dactylidis* | NRRL 29298 | KM361654 | KM361672 | - |
| *Fusarium dactylidis* | NRRL 29380 | KM361653 | KM361671 | - |
| *Fusarium decemcellulare* | KNU01 | LC212975 | LC214751 | - |
| *Fusarium decemcellulare* | NRRL 13412 | JX171567 | JX171567 | - |
| *Fusarium delphinoides* | NRRL 36160 | HM347204 | HM347219 | - |
| *Fusarium denticulatum* | CBS 407.97 | MT010953 | MT010970 | MT011002 |
| *Fusarium dimerum* | NRRL 20691 | JX171478 | JX171592 | - |
| *Fusarium dimerum* | NRRL 36140 | HM347203 | HM347218 | - |
| *Fusarium dlamini* | NRRL 13164 | KU171681 | KU171701 | - |
| *Fusarium domesticum* | NRRL 29976 | JX171528 | JX171641 | - |
| *Fusarium duoseptatum* | FocMa43 (VCG01217) | - | LS479207 | LS479653 |
| *Fusarium duoseptatum* | InaCC F828 | LS479520 | LS479266 | LS479715 |
| *Fusarium duoseptatum* | InaCC F829 | LS479528 | LS479274 | LS479723 |
| *Fusarium duoseptatum* | InaCC F831 | LS479538 | LS479285 | LS479734 |
| *Fusarium duoseptatum* | InaCC F835 | LS479567 | LS479315 | LS479764 |
| *Fusarium duoseptatum* | InaCC F911 | - | LS479234 | LS479683 |
| *Fusarium duoseptatum* | InaCC F915 | LS479494 | LS479238 | LS479687 |
| *Fusarium duoseptatum* | InaCC F916 | LS479495 | LS479239 | LS479688 |
| *Fusarium duoseptatum* | InaCC F920 | LS479499 | LS479244 | LS479693 |
| *Fusarium duoseptatum* | InaCC F921 | LS479500 | LS479245 | LS479694 |
| *Fusarium duoseptatum* | InaCC F975 | LS479549 | LS479296 | LS479745 |
| *Fusarium duoseptatum* | InaCC F976 | LS479550 | LS479297 | LS479746 |
| *Fusarium duoseptatum* | InaCC F977 | LS479551 | LS479298 | LS479747 |
| *Fusarium duoseptatum* | InaCC F978 | LS479552 | LS479299 | LS479748 |
| *Fusarium duoseptatum* | InaCC F979 | LS479553 | LS479300 | LS479749 |
| *Fusarium duoseptatum* | Indo80 | LS479619 | LS479387 | LS479829 |
| *Fusarium duoseptatum* | NRRL 36115 (VCG01224) | LS479475 | LS479218 | LS479666 |
| *Fusarium duoseptatum* | NRRL 36116 (VCG01223) | - | LS479219 | LS479667 |
| *Fusarium duoseptatum* * | Foc 38 | - | XX000000 | XX000000 |
| *Fusarium ensiforme* | CPC 27190 | - | LT746312 | LT746199 |
| *Fusarium ensiforme* | CPC 27191 | - | LT746313 | LT746200 |
| *Fusarium equiseti* | FUS18 | MN692709 | MN692731 | - |
| *Fusarium equiseti* | FUS28 | MN692714 | MN692736 | - |
| *Fusarium equiseti* | FUS42 | MN692718 | MN692740 | - |
| *Fusarium equiseti* | ITEM 10675 | - | LN901607 | LN901573 |
| *Fusarium equiseti* | ITEM 11363 | - | LN901609 | LN901574 |
| *Fusarium equiseti* | NRRL 43636 | HM347189 | GQ505841 | - |
| *Fusarium euwallaceae* | NRRL 54724 | JQ038023 | JQ038030 | - |
| *Fusarium euwallaceae* | NRRL 54725 | JQ038024 | JQ038031 | - |
| *Fusarium euwallaceae* | NRRL 54726 | JQ038025 | JQ038032 | - |
| *Fusarium falciforme* | MIW 58 | MN242937 | MN725019 | - |
| *Fusarium falciforme* | NRRL 43529 | JX171541 | JX171653 | - |
| *Fusarium falciforme* | NRRL 43529 | JX171541 | JX171653 | - |
| *Fusarium ficicrescens* | CBS 125178 | MT010950 | KT154002 | MT011004 |
| *Fusarium flagelliforme* | ITEM 11296 | - | LN901606 | LN901572 |
| *Fusarium flocciferum* | NRRL 25473 | JX171514 | JX171627 | - |
| *Fusarium flocciferum* | NRRL 45999 | HM347195 | GQ505497 | - |
| *Fusarium floridanum* | NRRL 62608 | KC691592 | KC691623 | - |
| *Fusarium floridanum* | NRRL 62628 | KC691593 | KC691624 | - |
| *Fusarium floridanum* | NRRL 62629 | KC691594 | KC691625 | - |
| *Fusarium foetens* | CBS 110286 | MT010945 | MT010984 | MT011001 |
| *Fusarium foetens* | NRRL 38302 | JX171652 | JX171652 | - |
| *Fusarium fujikuroi* | NRRL 13566 | JX171456 | JX171570 | - |
| *Fusarium fujikuroi* | NRRL 5538 | MN193916 | MN193888 | - |
| *Fusarium fujikuroi* | NRRL 66288 | MG282385 | MG282415 | - |
| *Fusarium gaditjirrii* | NRRL 45417 | JX171654 | JX171654 | - |
| *Fusarium globosum* | NRRL 26132 | LT746301 | LT746343 | - |
| *Fusarium globosum* | NRRL 26133 | LT746302 | LT746344 | - |
| *Fusarium globosum* | NRRL 26134 | LT746303 | LT746345 | - |
| *Fusarium gracilipes* | NRRL 43635 | HM347188 | GQ505840 | - |
| *Fusarium graminearum* | SP100 | - | MN625698 | MK611901 |
| *Fusarium graminearum* | SP102 | - | MN625699 | MK611900 |
| *Fusarium graminearum* | SP99 | - | MN625697 | MK611899 |
| *Fusarium graminum* | NRRL 20692 | JX171479 | JX171593 | - |
| *Fusarium grosmichelii* | InaCC F832 | LS479542 | LS479289 | LS479738 |
| *Fusarium grosmichelii* | InaCC F833 | LS479548 | LS479295 | LS479744 |
| *Fusarium grosmichelii* | InaCC F848 | LS479588 | LS479338 | LS479786 |
| *Fusarium grosmichelii* | InaCC F849 | LS479589 | LS479339 | LS479787 |
| *Fusarium grosmichelii* | InaCC F850 | - | LS479340 | LS479788 |
| *Fusarium grosmichelii* | InaCC F851 | - | LS479341 | LS479789 |
| *Fusarium grosmichelii* | InaCC F852 | - | LS479342 | LS479790 |
| *Fusarium grosmichelii* | InaCC F853 | - | LS479343 | LS479791 |
| *Fusarium grosmichelii* | InaCC F854 | LS479591 | LS479345 | LS479793 |
| *Fusarium grosmichelii* | InaCC F855 | LS479592 | LS479346 | LS479794 |
| *Fusarium grosmichelii* | InaCC F861 | LS479597 | LS479351 | LS479797 |
| *Fusarium grosmichelii* | InaCC F862 | LS479598 | LS479352 | LS479798 |
| *Fusarium grosmichelii* | InaCC F863 | LS479599 | LS479353 | LS479799 |
| *Fusarium grosmichelii* | InaCC F867 | - | LS479360 | LS479806 |
| *Fusarium grosmichelii* | InaCC F868 | - | LS479361 | LS479807 |
| *Fusarium grosmichelii* | InaCC F884 | LS479616 | LS479382 | LS479824 |
| *Fusarium grosmichelii* | InaCC F887 | LS479620 | LS479388 | LS479830 |
| *Fusarium grosmichelii* | InaCC F888 | LS479621 | LS479389 | LS479831 |
| *Fusarium grosmichelii* | Indo83 | - | LS479390 | - |
| *Fusarium grosmichelii* | NRRL 36120 (VCG01218) | LS479478 | LS479222 | LS479670 |
| *Fusarium guilinense* | NRRL 32865 | HM347161 | GQ505792 | - |
| *Fusarium guttiforme* | CBS 409.97 | MT010938 | MT010967 | MT010999 |
| *Fusarium guttiforme* | NRRL 2294 | MN193917 | MN193889 | - |
| *Fusarium guttiforme* | NRRL 22945 | JX171505 | JX171618 | - |
| *Fusarium hainanense* | NRRL 26417 | JX171522 | GQ505776 | - |
| *Fusarium heterosporum* | NRRL 20693 | JX171480 | JX171594 | - |
| *Fusarium hexaseptatum* | InaCC F866 | - | LS479359 | LS479805 |
| *Fusarium hostae* | NRRL 29888 | MT409435 | MT409445 | - |
| *Fusarium hostae* | NRRL 29889 | JX171640 | JX171640 | - |
| *Fusarium humicola* | CBS 124.73 | MN120718 | MN120738 | - |
| *Fusarium illudens* | NRRL 22090 | JX171488 | JX171601 | - |
| *Fusarium incarnatum* | ITEM 6748 | - | LN901618 | LN901582 |
| *Fusarium incarnatum* | ITEM 7155 | - | LN901617 | LN901581 |
| *Fusarium incarnatum* | NRRL 32866 | HM347162 | GQ505793 | - |
| *Fusarium ipomoeae* | NRRL 43640 | HM347191 | GQ505845 | - |
| *Fusarium irregulare* | NRRL 32175 | JX171532 | GQ505787 | - |
| *Fusarium irregulare* | NRRL 34006 | HM347169 | GQ505808 | - |
| *Fusarium kalimantanense* | InaCC F917 | LS479497 | LS479241 | LS479690 |
| *Fusarium kalimantanense* | InaCC F918 | - | LS479242 | LS479691 |
| *Fusarium kalimantanense* | InaCC F922 | - | LS479246 | LS479695 |
| *Fusarium keratoplasticum* | ATS92 | MF179712 | KY512536 | - |
| *Fusarium keratoplasticum* | ATS94 | MF179714 | KY512538 | - |
| *Fusarium kyushuense* | NRRL 66296 | MG282364 | MG282393 | - |
| *Fusarium lacertarum* | NRRL 20423 | JX171581 | GQ505771 | - |
| *Fusarium lactis* | CBS 411.97 | MT010954 | MT010969 | MT011010 |
| *Fusarium langsethiae* | NRRL 54940 | JX171550 | JX171662 | - |
| *Fusarium lateritium* | NRRL 13622 | JX171457 | HM068350 | - |
| *Fusarium lateritium* | NRRL 25197 | HM347140 | HM347207 | - |
| *Fusarium longipes* | NRRL 13368 | JX171448 | JX171562 | - |
| *Fusarium longipes* | NRRL 13374 | JX171450 | JX171564 | - |
| *Fusarium longipes* | NRRL 20723 | JX171483 | JX171596 | - |
| *Fusarium luffae* | NRRL 32522 | HM347158 | GQ505790 | - |
| *Fusarium lunatum* | NRRL 36168 | JX171536 | JX171648 | - |
| *Fusarium lunulosporum* | NRRL 13393 | KM361637 | KM361655 | - |
| *Fusarium lyarnte* | NRRL 54252 | JX171661 | MN193908 | - |
| *Fusarium macrosporum* | CPC 28191 | - | LT746331 | LT746218 |
| *Fusarium macrosporum* | CPC 28192 | - | LT746332 | LT746219 |
| *Fusarium macrosporum* | CPC 28193 | - | LT746333 | LT746220 |
| *Fusarium mangiferae* | MUCL 54671 | - | LT575059 | LT574978 |
| *Fusarium mangiferae* | NRRL 25226 | JX171509 | HM068353 | - |
| *Fusarium mangiferae* | UMAF 910 | KP753434 | KP753441 | - |
| *Fusarium meridionale* | NRRL 28436 | KM361642 | KM361660 | - |
| *Fusarium mesoamericanum* | NRRL 25797 | KM361639 | KM361657 | - |
| *Fusarium mexicanum* | MICMW 32.13a | MN242900 | MN724975 | - |
| *Fusarium mexicanum* | MICMW 3A | MN242905 | MN724980 | - |
| *Fusarium miscanthi* | NRRL 26231 | JX171634 | JX171634 | - |
| *Fusarium multiceps* | NRRL 43639 | HM347190 | GQ505844 | - |
| *Fusarium mundagurra* | RBG5717 | KP083272 | KP083276 | - |
| *Fusarium musae* | CBS 624.87 | MT010957 | MT010973 | MT010991 |
| *Fusarium nanum* | NRRL 32868 | HM347163 | GQ505795 | - |
| *Fusarium napiforme* | CBS 748.97 | MT010958 | KU604233 | MT011011 |
| *Fusarium napiforme* | F111 | KU974338 | KU974364 | - |
| *Fusarium napiforme* | NRRL 25196 | MN193919 | MN193891 | - |
| *Fusarium nectrioides* | NRRL 20689 | JX171477 | JX171591 | - |
| *Fusarium nelsonii* | NRRL 13338 | JX171447 | GQ505466 | - |
| *Fusarium nematophilum* | NRRL 54600 | JX171552 | JX171664 | - |
| *Fusarium neocosmosporiellum* | NRRL 22436 | JX171497 | JX171610 | - |
| *Fusarium newnesense* | RBG5443 | KJ397218 | KJ397254 | - |
| *Fusarium nisikadoi* | NRRL 25179 | JX171620 | JX171620 | - |
| *Fusarium nisikadoi* | NRRL 25203 | MG282388 | MG282418 | - |
| *Fusarium nisikadoi* | NRRL 25308 | MG282391 | MG282421 | - |
| *Fusarium nodosum* | CBS 200.63 | MN120724 | MN120742 | - |
| *Fusarium nodosum* | CBS 201.63 | MN120725 | MN120743 | - |
| *Fusarium nodosum* | CBS 698.74 | MN120726 | MN120744 | - |
| *Fusarium nurragi* | NRRL 36452 | JX171538 | JX171650 | - |
| *Fusarium nygamai* | CBS 749.97 | MT010955 | KU604262 | MT011009 |
| *Fusarium nygamai* | NRRL 66291 | MG282368 | MG282397 | - |
| *Fusarium nygamai* | NRRL 66293 | MG282367 | MG282396 | - |
| *Fusarium obliquiseptatum* | NRRL 62610 | KC691605 | KC691636 | - |
| *Fusarium obliquiseptatum* | NRRL 62611 | KC691606 | KC691637 | - |
| *Fusarium odoratissimum* | InaCC F1000 | LS479575 | LS479323 | LS479772 |
| *Fusarium odoratissimum* | InaCC F816 | LS479485 | LS479228 | LS479677 |
| *Fusarium odoratissimum* | InaCC F817 | LS479556 | LS479304 | LS479753 |
| *Fusarium odoratissimum* | InaCC F818 | LS479584 | LS479333 | LS479782 |
| *Fusarium odoratissimum* | InaCC F819 | LS479600 | LS479354 | LS479800 |
| *Fusarium odoratissimum* | InaCC F821 | LS479609 | LS479374 | LS479818 |
| *Fusarium odoratissimum* | InaCC F822 | LS479618 | LS479386 | LS479828 |
| *Fusarium odoratissimum* | InaCC F824 | LS479486 | LS479229 | LS479678 |
| *Fusarium odoratissimum* | InaCC F825 | LS479496 | LS479240 | LS479689 |
| *Fusarium odoratissimum* | InaCC F836 | LS479577 | LS479325 | LS479774 |
| *Fusarium odoratissimum* | InaCC F837 | LS479578 | LS479326 | LS479775 |
| *Fusarium odoratissimum* | InaCC F838 | LS479579 | LS479327 | LS479776 |
| *Fusarium odoratissimum* | InaCC F839 Indo25 (VCG01219) | LS479580 | LS479328 | LS479777 |
| *Fusarium odoratissimum* | InaCC F840 | - | LS479329 | LS479778 |
| *Fusarium odoratissimum* | InaCC F846 | - | LS479336 | LS479785 |
| *Fusarium odoratissimum* | InaCC F857 | LS479594 | LS479348 | LS479795 |
| *Fusarium odoratissimum* | InaCC F864 | - | LS479356 | LS479802 |
| *Fusarium odoratissimum* | InaCC F870 | LS479602 | LS479363 | LS479809 |
| *Fusarium odoratissimum* | InaCC F871 | - | LS479365 | LS479811 |
| *Fusarium odoratissimum* | InaCC F873 | LS479604 | LS479369 | LS479814 |
| *Fusarium odoratissimum* | InaCC F875 | LS479607 | LS479372 | LS479816 |
| *Fusarium odoratissimum* | InaCC F876 | LS479608 | LS479373 | LS479817 |
| *Fusarium odoratissimum* | InaCC F877 | LS479610 | LS479375 | LS479819 |
| *Fusarium odoratissimum* | InaCC F879 | LS479612 | LS479377 | LS479820 |
| *Fusarium odoratissimum* | InaCC F880 | - | LS479378 | LS479821 |
| *Fusarium odoratissimum* | InaCC F882 | LS479614 | LS479380 | LS479822 |
| *Fusarium odoratissimum* | InaCC F883 | LS479615 | LS479381 | LS479823 |
| *Fusarium odoratissimum* | InaCC F885 | - | LS479384 | LS479826 |
| *Fusarium odoratissimum* | InaCC F891 | - | LS479393 | LS479833 |
| *Fusarium odoratissimum* | InaCC F892 | LS479624 | LS479394 | LS479834 |
| *Fusarium odoratissimum* | InaCC F893 | LS479625 | LS479395 | LS479835 |
| *Fusarium odoratissimum* | InaCC F894 | LS479626 | LS479396 | LS479836 |
| *Fusarium odoratissimum* | InaCC F896 | LS479629 | LS479399 | LS479839 |
| *Fusarium odoratissimum* | InaCC F897 | LS479630 | LS479400 | LS479840 |
| *Fusarium odoratissimum* | InaCC F898 | LS479631 | LS479401 | LS479841 |
| *Fusarium odoratissimum* | InaCC F899 | LS479632 | LS479402 | LS479842 |
| *Fusarium odoratissimum* | InaCC F900 | LS479633 | LS479403 | LS479843 |
| *Fusarium odoratissimum* | InaCC F901 | LS479634 | LS479404 | LS479844 |
| *Fusarium odoratissimum* | InaCC F902 | LS479635 | LS479405 | LS479845 |
| *Fusarium odoratissimum* | InaCC F903 | LS479636 | LS479406 | LS479846 |
| *Fusarium odoratissimum* | InaCC F904 | LS479637 | LS479407 | LS479847 |
| *Fusarium odoratissimum* | InaCC F905 | LS479638 | LS479408 | LS479848 |
| *Fusarium odoratissimum* | InaCC F906 | LS479639 | LS479409 | LS479849 |
| *Fusarium odoratissimum* | InaCC F907 | LS479487 | LS479230 | LS479679 |
| *Fusarium odoratissimum* | InaCC F908 | LS479488 | LS479231 | LS479680 |
| *Fusarium odoratissimum* | InaCC F909 | LS479489 | LS479232 | LS479681 |
| *Fusarium odoratissimum* | InaCC F910 | LS479490 | LS479233 | LS479682 |
| *Fusarium odoratissimum* | InaCC F912 | LS479491 | LS479235 | LS479684 |
| *Fusarium odoratissimum* | InaCC F919 | LS479498 | LS479243 | LS479692 |
| *Fusarium odoratissimum* | InaCC F923 | LS479501 | LS479247 | LS479696 |
| *Fusarium odoratissimum* | InaCC F924 | LS479502 | LS479248 | LS479697 |
| *Fusarium odoratissimum* | InaCC F925 | LS479503 | LS479249 | LS479698 |
| *Fusarium odoratissimum* | InaCC F926 | LS479504 | LS479250 | LS479699 |
| *Fusarium odoratissimum* | InaCC F927 | LS479506 | LS479252 | LS479701 |
| *Fusarium odoratissimum* | InaCC F928 | LS479507 | LS479253 | LS479702 |
| *Fusarium odoratissimum* | InaCC F929 | LS479508 | LS479254 | LS479703 |
| *Fusarium odoratissimum* | InaCC F930 | LS479509 | LS479255 | LS479704 |
| *Fusarium odoratissimum* | InaCC F931 | LS479510 | LS479256 | LS479705 |
| *Fusarium odoratissimum* | InaCC F932 | LS479511 | LS479257 | LS479706 |
| *Fusarium odoratissimum* | InaCC F933 | LS479512 | LS479258 | LS479707 |
| *Fusarium odoratissimum* | InaCC F934 | LS479514 | LS479260 | LS479709 |
| *Fusarium odoratissimum* | InaCC F935 | LS479515 | LS479261 | LS479710 |
| *Fusarium odoratissimum* | InaCC F936 | LS479516 | LS479262 | LS479711 |
| *Fusarium odoratissimum* | InaCC F937 | LS479517 | LS479263 | LS479712 |
| *Fusarium odoratissimum* | InaCC F938 | LS479518 | LS479264 | LS479713 |
| *Fusarium odoratissimum* | InaCC F939 | LS479519 | LS479265 | LS479714 |
| *Fusarium odoratissimum* | InaCC F942 | LS479521 | LS479267 | LS479716 |
| *Fusarium odoratissimum* | InaCC F943 | LS479522 | LS479268 | LS479717 |
| *Fusarium odoratissimum* | InaCC F944 | LS479523 | LS479269 | LS479718 |
| *Fusarium odoratissimum* | InaCC F945 | LS479524 | LS479270 | LS479719 |
| *Fusarium odoratissimum* | InaCC F946 | LS479525 | LS479271 | LS479720 |
| *Fusarium odoratissimum* | InaCC F947 | LS479526 | LS479272 | LS479721 |
| *Fusarium odoratissimum* | InaCC F948 | LS479527 | LS479273 | LS479722 |
| *Fusarium odoratissimum* | InaCC F953 | LS479529 | LS479275 | LS479724 |
| *Fusarium odoratissimum* | InaCC F954 | LS479530 | LS479276 | LS479725 |
| *Fusarium odoratissimum* | InaCC F955 | LS479531 | LS479277 | LS479726 |
| *Fusarium odoratissimum* | InaCC F973 | LS479547 | LS479294 | LS479743 |
| *Fusarium odoratissimum* | InaCC F985 | LS479562 | LS479310 | LS479759 |
| *Fusarium odoratissimum* | InaCC F986 | LS479563 | LS479311 | LS479760 |
| *Fusarium odoratissimum* | InaCC F989 | LS479566 | LS479314 | LS479763 |
| *Fusarium odoratissimum* | InaCC F990 | LS479568 | LS479316 | LS479765 |
| *Fusarium odoratissimum* | InaCC F994 | LS479569 | LS479317 | LS479766 |
| *Fusarium odoratissimum* | InaCC F997 | LS479572 | LS479320 | LS479769 |
| *Fusarium odoratissimum* | InaCC F998 | LS479573 | LS479321 | LS479770 |
| *Fusarium odoratissimum* | InaCC F999 | LS479574 | LS479322 | LS479771 |
| *Fusarium odoratissimum* | Indo204 | LS479561 | LS479309 | LS479758 |
| *Fusarium odoratissimum* | Indo222 | LS479576 | LS479324 | LS479773 |
| *Fusarium odoratissimum* | Indo4 | LS479590 | LS479344 | LS479792 |
| *Fusarium odoratissimum* | Indo53 | - | LS479357 | LS479803 |
| *Fusarium odoratissimum* | Indo61 | - | LS479366 | LS479812 |
| *Fusarium odoratissimum* | Indo62 | - | LS479367 | - |
| *Fusarium odoratissimum* | Indo66 | LS479605 | LS479370 | LS479815 |
| *Fusarium odoratissimum* | Indo77 | LS479617 | LS479383 | LS479825 |
| *Fusarium odoratissimum* | Indo89 | LS479627 | LS479397 | LS479837 |
| *Fusarium odoratissimum* | JV11 | LS479465 | LS479205 | LS479651 |
| *Fusarium odoratissimum* | Leb1.2C | LS479466 | LS479206 | LS479652 |
| *Fusarium odoratissimum* | NRRL 36102 (VCG0121) | LS479468 | LS479209 | LS479655 |
| *Fusarium odoratissimum* | Pak1.1A | LS479479 | LS479223 | LS479671 |
| *Fusarium odoratissimum* * | Foc 56 | XX000000 | XX000000 | XX000000 |
| *Fusarium odoratissimum* * | Foc 61 | - | XX000000 | XX000000 |
| *Fusarium odoratissimum* | FocII5 (VCG01213) | LS479459 | LS479198 | LS479644 |
| *Fusarium penzigii* | NRRL 20711 | HM347202 | HM347217 | - |
| *Fusarium pernambucanum* | NRRL 32864 | HM347160 | GQ505791 | - |
| *Fusarium perseae* | CPC 26829 | - | LT991909 | LT991902 |
| *Fusarium perseae* | CPC 26830 | - | LT991910 | LT991903 |
| *Fusarium perseae* | CPC 26832 | - | LT991912 | LT991905 |
| *Fusarium peruvianum* | CBS 511.75 | MN120728 | MN120746 | - |
| *Fusarium petersiae* | JW14004 | MG386138 | MG386149 | - |
| *Fusarium petersiae* | JW14005 | MG386139 | MG386150 | - |
| *Fusarium phaseoli* | CBS 265.50 | KM232226 | KM232375 | HE647964 |
| *Fusarium phaseoli* | NRRL 22276 | JX171495 | JX171608 | - |
| *Fusarium phaseoli* | NRRL 22411 | KJ511267 | KJ511278 | - |
| *Fusarium phialophorum* | FocIndo25 | LS479464 | LS479204 | LS479650 |
| *Fusarium phialophorum* | FocST4.98 (VCG0120) | LS479484 | LS479227 | LS479676 |
| *Fusarium phialophorum* | InaCC F826 | LS479505 | LS479251 | LS479700 |
| *Fusarium phialophorum* | InaCC F827 | LS479513 | LS479259 | LS479708 |
| *Fusarium phialophorum* | InaCC F830 | LS479536 | LS479282 | LS479731 |
| *Fusarium phialophorum* | InaCC F834 | LS479557 | LS479305 | LS479754 |
| *Fusarium phialophorum* | InaCC F842 | LS479582 | LS479331 | LS479780 |
| *Fusarium phialophorum* | InaCC F843 | LS479583 | LS479332 | LS479781 |
| *Fusarium phialophorum* | InaCC F844 | LS479585 | LS479334 | LS479783 |
| *Fusarium phialophorum* | InaCC F845 | LS479586 | LS479335 | LS479784 |
| *Fusarium phialophorum* | InaCC F889 Indo84 (VCG01216) | LS479622 | LS479391 | LS479832 |
| *Fusarium phialophorum* | InaCC F969 | LS479543 | LS479290 | LS479739 |
| *Fusarium phialophorum* | InaCC F970 | LS479544 | LS479291 | LS479740 |
| *Fusarium phialophorum* | InaCC F971 | LS479545 | LS479292 | LS479741 |
| *Fusarium phialophorum* | InaCC F972 | LS479546 | LS479293 | LS479742 |
| *Fusarium phialophorum* | InaCC F981 | - | LS479303 | LS479752 |
| *Fusarium phialophorum* | InaCC F982 | LS479558 | LS479306 | LS479755 |
| *Fusarium phialophorum* | InaCC F987 | LS479564 | LS479312 | LS479761 |
| *Fusarium phialophorum* | InaCC F995 | LS479570 | LS479318 | LS479767 |
| *Fusarium phialophorum* | InaCC F996 | LS479571 | LS479319 | LS479768 |
| *Fusarium phialophorum* | NRRL 36101 (VCG0123) | LS479467 | LS479208 | LS479654 |
| *Fusarium phialophorum* | NRRL 36103 (VCG0122) | LS479469 | LS479210 | LS479656 |
| *Fusarium phialophorum* | NRRL 36109 (VCG01211) | LS479471 | LS479214 | LS479661 |
| *Fusarium phialophorum* | NRRL 36112 (VCG01215) | LS479473 | LS479216 | LS479664 |
| *Fusarium phialophorum* | R1.0124 | LS479483 | - | LS479675 |
| *Fusarium phyllophilum* | NRRL 13617 | KF466399 | KF466410 | - |
| *Fusarium plagianthi* | NRRL 22632 | JX171501 | JX171614 | - |
| *Fusarium poae* | NRRL 13714 | JX171458 | JX171572 | - |
| *Fusarium poae* | NRRL 66297 | MG282363 | MG282392 | - |
| *Fusarium praegraminearum* | NRRL 39664 | KX260125 | KX260126 | - |
| *Fusarium proliferatum* | ITEM2287 | LT841251 | LT841252 | LT841245 |
| *Fusarium proliferatum* | ITEM2400 | LT841265 | LT841266 | LT841259 |
| *Fusarium proliferatum* | NRRL 22944 | JX171504 | HM068352 | - |
| *Fusarium pseudensiforme* | NRRL 46517 | KC691615 | KC691645 | - |
| *Fusarium pseudoanthophilum* | CBS 414.97 | MT010949 | MT010980 | MT011006 |
| *Fusarium pseudocircinatum* | NRRL 22946 | MG838070 | MN724939 | - |
| *Fusarium pseudocircinatum* | NRRL 31631 | MG838073 | MN724942 | - |
| *Fusarium pseudocircinatum* | NRRL 53570 | MG838075 | MN724944 | - |
| *Fusarium pseudograminearum* | NRRL 28062 | JX171524 | JX171637 | - |
| *Fusarium pseudograminearum* | NRRL 28065 | MG282389 | MG282419 | - |
| *Fusarium pseudonygamai* | CBS 417.97 | MT010951 | MT010978 | MT011008 |
| *Fusarium purpurascens* | InaCC F823 | LS479628 | LS479398 | LS479838 |
| *Fusarium purpurascens* | InaCC F886 | - | LS479385 | LS479827 |
| *Fusarium purpurascens* | InaCC F913 | LS479492 | LS479236 | LS479685 |
| *Fusarium purpurascens* | InaCC F914 | LS479493 | LS479237 | LS479686 |
| *Fusarium purpurascens* | InaCC F966 | LS479539 | LS479286 | LS479735 |
| *Fusarium purpurascens* | InaCC F967 | LS479540 | LS479287 | LS479736 |
| *Fusarium purpurascens* | InaCC F968 | LS479541 | LS479288 | LS479737 |
| *Fusarium purpurascens* | NRRL 36107 (VCG0126) | - | LS479213 | LS479659 |
| *Fusarium ramigenum* | CBS 418.97 | MT010959 | MT010975 | MT011012 |
| *Fusarium ramigenum* | NRRL 25208 | KF466401 | KF466412 | - |
| *Fusarium redolens* | CBS 743.97 | MT010935 | MT010961 | MT010987 |
| *Fusarium redolens* | NRRL 22901 | JX171616 | JX171616 | - |
| *Fusarium redolens* | NRRL 25600 | MT409433 | MT409443 | - |
| *Fusarium ruscii* | NRRL 22134 | JX171490 | JX171603 | - |
| *Fusarium sacchari* | CBS 147.25 | MT010941 | MT010962 | MT010988 |
| *Fusarium sacchari* | NRRL 44901 | HM347194 | HM347212 | - |
| *Fusarium sacchari* | YN BS37 | MK983434 | MK829737 | - |
| *Fusarium salinense* | CPC 26403 | LT746284 | LT746304 | LT746191 |
| *Fusarium salinense* | CPC 26457 | LT746285 | LT746305 | LT746192 |
| *Fusarium salinense* | CPC 26973 | LT746286 | LT746306 | LT746193 |
| *Fusarium sambucinum* | NRRL 22187 | JX171493 | JX171606 | - |
| *Fusarium sangayamense* | InaCC F961 | - | LS479284 | LS479733 |
| *Fusarium sarcochroum* | CPC 28075 | LT746296 | LT746324 | LT746211 |
| *Fusarium sarcochroum* | CPC 28116 | LT746297 | LT746325 | LT746212 |
| *Fusarium sarcochroum* | NRRL 20472 | JX171472 | JX171586 | - |
| *Fusarium scirpi* | NRRL 13402 | JX171452 | GQ505770 | - |
| *Fusarium setosum* | NRRL 36526 | JX171539 | JX171651 | - |
| *Fusarium siculi* | CPC 27188 | LT746299 | LT746327 | LT746214 |
| *Fusarium siculi* | CPC 27189 | LT746300 | LT746328 | LT746215 |
| *Fusarium solani* | FBF7 | - | MK606410 | MK606409 |
| *Fusarium solani* | LEMM_110148 | - | LN828050 | LN827961 |
| *Fusarium solani* | LEMM_110266 | - | LN828053 | LN827964 |
| *Fusarium solani melongenae* | MIM 28 | MN242938 | MN725021 | - |
| *Fusarium solani melongenae* | MIW 81 | MN242939 | MN724934 | - |
| *Fusarium solani melongenae* | NRRL 22147 | MG282390 | MG282420 | - |
| *Fusarium spinosum* | CBS 122438 | MN120729 | MN120747 | - |
| *Fusarium spinosum* | NRRL 43631 | HM347187 | GQ505491 | - |
| *Fusarium sporodochiale* | CBS 199.63 | MN120730 | MN120748 | - |
| *Fusarium sporodochiale* | CBS 220.61 | MN120731 | MN120749 | - |
| *Fusarium sporotrichioides* | NRRL 25479 | HM347144 | HM347210 | - |
| *Fusarium sporotrichioides* | NRRL 3299 | JX171444 | DQ676587 | - |
| *Fusarium sporotrichioides* | NRRL 66295 | MG282378 | MG282408 | - |
| *Fusarium staphyleae* | NRRL 22316 | JX171496 | JX171609 | - |
| *Fusarium sterilihyphosum* | NRRL 25623 | LR792581 | LR792617 | - |
| *Fusarium stilboides* | HA 1 2 | MK887361 | MK887362 | - |
| *Fusarium stilboides* | NRRL 20429 | JX171468 | JX171582 | - |
| *Fusarium subglutinans* | NRRL 22016 | JX171486 | JX171599 | - |
| *Fusarium subglutinans* | NRRL 54158 | HM347201 | HM347216 | - |
| *Fusarium subglutinans* | NRRL 66333 | MN193926 | MN193898 | - |
| *Fusarium sublunatum* | NRRL 13384 | JX171451 | JX171565 | - |
| *Fusarium subtropicale* | NRRL 66764 | MH706972 | MH706973 | - |
| *Fusarium sulawesiense* | NRRL 34004 | HM347167 | GQ505806 | - |
| *Fusarium tanahbumbuense* | CBS 101138 | MN120733 | MN120751 | - |
| *Fusarium tanahbumbuense* | NRRL 34005 | HM347168 | GQ505807 | - |
| *Fusarium tardichlamydosporum* | BRIP44611 | KX434927 | KX434962 | - |
| *Fusarium tardichlamydosporum* | BRIP62955 | KX434936 | KX434971 | - |
| *Fusarium tardichlamydosporum* * | Foc 1 | - | XX000000 | XX000000 |
| *Fusarium tardichlamydosporum* * | Foc 11 | - | XX000000 | XX000000 |
| *Fusarium tardichlamydosporum* * | Foc 16 | - | XX000000 | XX000000 |
| *Fusarium tardichlamydosporum* * | Foc 18 | - | XX000000 | XX000000 |
| *Fusarium tardichlamydosporum* * | Foc 2 | - | XX000000 | XX000000 |
| *Fusarium tardichlamydosporum* * | Foc 21 | - | XX000000 | XX000000 |
| *Fusarium tardichlamydosporum* * | Foc 23-2 | - | XX000000 | XX000000 |
| *Fusarium tardichlamydosporum* * | Foc 24 | XX000000 | XX000000 | XX000000 |
| *Fusarium tardichlamydosporum* * | Foc 25-1 | - | XX000000 | XX000000 |
| *Fusarium tardichlamydosporum* * | Foc 25-2 | - | XX000000 | XX000000 |
| *Fusarium tardichlamydosporum* * | Foc 5 | - | XX000000 | XX000000 |
| *Fusarium tardichlamydosporum* * | Foc 6 | - | XX000000 | XX000000 |
| *Fusarium tardichlamydosporum* * | Foc 7 | - | XX000000 | XX000000 |
| *Fusarium tardichlamydosporum* | InaCC F956 | LS479532 | LS479278 | LS479727 |
| *Fusarium tardichlamydosporum* | InaCC F957 | LS479533 | LS479279 | LS479728 |
| *Fusarium tardichlamydosporum* | InaCC F958 | LS479534 | LS479280 | LS479729 |
| *Fusarium tardichlamydosporum* | InaCC F959 | LS479535 | LS479281 | LS479730 |
| *Fusarium tardichlamydosporum* | NRRL 36105 (VCG0124) | LS479470 | LS479211 | LS479657 |
| *Fusarium tardichlamydosporum* | NRRL 36106 (VCG0125) | - | LS479212 | LS479658 |
| *Fusarium tardichlamydosporum* | NRRL 36111 (VCG0128) | LS479472 | LS479215 | LS479663 |
| *Fusarium tardichlamydosporum* | NRRL 36117 (VCG01222) | LS479476 | LS479220 | LS479668 |
| *Fusarium tardicrescens* | NRRL 36113 (VCG01214) | LS479474 | LS479217 | LS479665 |
| *Fusarium tardicrescens* | NRRL 37622 | LS479463 | LS479203 | LS479649 |
| *Fusarium tardicrescens* | NRRL 54005 | LS479482 | LS479226 | LS479674 |
| *Fusarium tardicrescens* | NRRL 54008 | LS479481 | LS479225 | LS479673 |
| *Fusarium temperatum* | NRRL 25622 | LR792582 | LR792618 | - |
| *Fusarium thapsinum* | NRRL 22045 | JX171487 | JX171600 | - |
| *Fusarium thapsinum* | NRRL 22049 | KU171713 | MN193899 | - |
| *Fusarium tjaetaba* | RBG5361 | KP083267 | KP083275 | - |
| *Fusarium tonkinense* | CPC 27195 | - | LT746340 | LT746227 |
| *Fusarium torreyae* | NRRL 54149 | JX171548 | HM068359 | - |
| *Fusarium torulosum* | NRRL 22748 | JX171502 | JX171615 | - |
| *Fusarium torulosum* | NRRL 52772 | JF741003 | MH582377 | - |
| *Fusarium tricinctum* | NRRL 25481 | JX171516 | HM068327 | - |
| *Fusarium tuaranense* | NRRL 22231 | KC691600 | KC691631 | - |
| *Fusarium tuaranense* | NRRL 46518 | KC691601 | KC691632 | - |
| *Fusarium tuaranense* | NRRL 46519 | KC691602 | KC691633 | - |
| *Fusarium tucumaniae* | NRRL 31086 | KJ511269 | KJ511280 | - |
| *Fusarium tucumaniae* | NRRL 34546 | KJ511273 | KJ511284 | - |
| *Fusarium tupiense* | NRRL 53984 | LR792583 | LR792619 | - |
| *Fusarium tupiense* | UMAF 0917 | KP753436 | KP753443 | - |
| *Fusarium tupiense* | UMAF 0933 | KP753437 | KP753444 | - |
| *Fusarium udum* | NRRL 25194 | MN193928 | MN193900 | - |
| *Fusarium ussurianum* | NRRL 45681 | KM361648 | KM361666 | - |
| *Fusarium vanettenii* | NRRL 45880 | JX171543 | JX171655 | - |
| *Fusarium venenatum* | NRRL 22196 | JX171494 | JX171607 | - |
| *Fusarium ventricosum* | NRRL 13953 | JX171461 | JX171575 | - |
| *Fusarium ventricosum* | NRRL 20846 | JX171484 | JX171597 | - |
| *Fusarium ventricosum* | NRRL 25729 | JX171520 | JX171633 | - |
| *Fusarium verticillioides* | YN DH24 | MK886821 | MK886821 | - |
| *Fusarium verticillioides* | YN DH28 | MK983460 | MK983384 | - |
| *Fusarium verticillioides* | YN SJ46 | MK983456 | MK983397 | - |
| *Fusarium virguliforme* | NRRL 31041 | JX171643 | FJ240386 | - |
| *Fusarium vorosii* | NRRL 37605 | KM361647 | KM361665 | - |
| *Fusarium xylarioides* | NRRL 25486 | JX171630 | HM068355 | - |
| *Fusarium xyrophilum* | NRRL 62710 | MN193931 | MN193903 | - |
| *Fusarium xyrophilum* | NRRL 62721 | MN193933 | MN193905 | - |
| *Fusarium xyrophilum* | NRRL 66890 | MN193932 | MN193904 | - |
| *Fusarium zealandicum* | NRRL 22465 | JX171498 | JX171611 | - |
| *Fusicolla aquaeductuum* | NRRL 20686 | JX171476 | JX171590 | - |
| *Fusicolla sp.* | NRRL 22136 | JX171491 | JX171604 | - |
| *Macroconia leptosphaeriae* | NRRL 54562 | JX171556 | JX171668 | - |
| *Macroconia sp.* | NRRL 54563 | JX171557 | JX171669 | - |
| *Microcera coccophila* | NRRL 13962 | JX171462 | JX171576 | - |
| *Microcera diploa* | NRRL 36545 | JX171463 | JX171577 | - |
| *Microcera larvarum* | NRRL 20473 | JX171473 | JX171587 | - |

## Notes

### Competing Interest Statement

The authors have declared no competing interest.

